# Non-random association of MHC-I alleles in favor of high diversity haplotypes in wild songbirds revealed by computer-assisted MHC haplotype inference using the R package MHCtools

**DOI:** 10.1101/2020.03.24.005207

**Authors:** Jacob Roved, Bengt Hansson, Martin Stervander, Dennis Hasselquist, Helena Westerdahl

## Abstract

Major histocompatibility complex (MHC) genes play a central role for pathogen recognition by the adaptive immune system. The MHC genes are often duplicated and tightly linked within a small genomic region. This structural organization suggests that natural selection acts on the combined property of multiple MHC gene copies in segregating haplotypes, rather than on single MHC genes. This may have important implications for analyses of patterns of selection on MHC genes. Here, we present a computer-assisted protocol to infer segregating MHC haplotypes from family data, based on functions in the R package MHCtools. We employed this method to identify 107 unique MHC class I (MHC-I) haplotypes in 116 families of wild great reed warblers (*Acrocephalus arundinaceus*). In our data, the MHC-I genes were tightly linked in haplotypes and inherited as single units, with only two observed recombination events among 334 offspring. We found substantial variation in the number of different MHC-I alleles per haplotype, and the divergence between alleles in MHC-I haplotypes was significantly higher than between randomly assigned alleles in simulated haplotypes. This suggests that selection has favored non-random associations of divergent MHC-I alleles in haplotypes to increase the range of pathogens that can be recognized by the adaptive immune system. Further studies of selection on MHC haplotypes in natural populations is an interesting avenue for future research. Moreover, inference and analysis of MHC haplotypes offers important insights into the structural organization of MHC genes, and may improve the accuracy of the MHC region in *de novo* genome assemblies.

## Introduction

The major histocompatibility complex (MHC) is a multigene family that plays a vital role in the vertebrate adaptive immune system (Klein & Sato, 2000). An ongoing arms race with pathogens has caused MHC genes to exhibit extreme levels of genetic diversity, and these genes have gathered a broad interest in studies of adaptive genetic variation in vertebrates (Ejsmond & Radwan, 2015; Kaufman, 2018; Piertney & Oliver, 2006). In humans, the MHC genes are found within a genomic region that spans approximately 4 Mb (Trowsdale & Knight, 2015). The MHC genes are often tightly linked and MHC alleles have been found to be non-randomly associated within haplotypes, suggesting that selection acts on multi-locus MHC haplotypes rather than single MHC genes (Begovich et al., 1992, 2001; Buhler, Nunes, & Sanchez-Mazas, 2016; Hollenbach et al., 2001; Kaufman, 1999; Testi et al., 2015). In MHC studies, the heterozygote advantage hypothesis is often used to describe the principle that individuals with two different maternally and paternally inherited MHC alleles should be able to recognize antigens from a larger range of pathogens, than individuals with two identical MHC alleles (Doherty & Zinkernagel, 1975; Hughes & Nei, 1992). However, from a functional perspective it should not matter whether the MHC molecules expressed in a cell are encoded by alleles that are harbored on the same or on different paternally or maternally inherited multi-locus haplotypes. This reasoning implies that haplotypes with a larger number of MHC gene copies may be favored by natural selection, because they confer an advantage in terms of presenting antigens from a larger range of pathogens, given that the genes have the same function and are expressed to the same degree. Though, note that negative selection of T-cells in the thymus is thought to associate too high individual MHC gene copy numbers with a disadvantage (Nowak, Tarczyhornoch, & Austyn, 1992; Woelfing, Traulsen, Milinski, & Boehm, 2009). Additionally, *in silico* models have shown that the degree of divergence between the amino acid sequences of MHC alleles is positively correlated with the combined number of different antigens that can be bound by the corresponding MHC molecules (Lenz, 2011; Pierini & Lenz, 2018). This suggests that haplotypes that combine highly divergent MHC alleles may be favored by natural selection in a manner similar to MHC heterozygotes (*i*.*e*., divergent allele advantage, sensu Wakeland et al. (1990)).

Detailed knowledge about MHC haplotype structure is mostly limited to humans and a few model organisms, but in recent years, there has been a growing interest in characterizing MHC haplotypes and investigating their effects also in wild non-model species (Gaigher et al., 2016, 2018; Huchard, Weill, Cowlishaw, Raymond, & Knapp, 2008; Niskanen et al., 2014; Sin et al., 2014). Gaigher et al. (2016) demonstrated how analysis of the segregation patterns of MHC alleles within families in a natural population of barn owls (*Tyto alba*) could be used to infer MHC haplotypes, and thereby obtain information about linkage and recombination, and confidently assess the number of MHC gene copies and the presence of gene copy number variation (Gaigher et al., 2016). In a follow-up study, they showed how MHC haplotype data can be employed to investigate non-random associations of MHC alleles in haplotypes in a wild animal population (Gaigher et al., 2018). The studies by Gaigher et al. demonstrate the significant value of family-assisted haplotype inference in future studies of MHC genes in evolutionary biology and ecology.

Since the advent of high throughput DNA sequencing, MHC genotyping in non-model organisms has mostly been carried out using PCR-based amplicon sequencing (Biedrzycka, Sebastian, Migalska, Westerdahl, & Radwan, 2017; Burri, Promerova, Goebel, & Fumagalli, 2014; Promerová et al., 2012; Zagalska-Neubauer et al., 2010). However, in many non-model species, amplification of specific MHC loci is impeded by the sequence similarity across different loci, caused by recombination and gene conversion within and between MHC genes, and in such cases it is often necessary to co-amplify multiple MHC loci (Alcaide, Liu, & Edwards, 2013; Burri et al., 2014; Zagalska-Neubauer et al., 2010). While this technique is useful for estimating the overall genetic diversity harbored in the MHC, the resulting data contain no information about linkage or spatial organization of the amplified alleles (Alcaide et al., 2013; Biedrzycka, Sebastian, et al., 2017; Burri et al., 2014; Gaigher et al., 2016). Furthermore, the number of loci has to be estimated indirectly from the number of different alleles detected in each sample, and associating alleles with specific loci becomes extremely difficult, in particular in species with highly duplicated MHC genes (Gaigher et al., 2016; Lighten, van Oosterhout, Paterson, Mcmullan, & Bentzen, 2014; O’Connor, Westerdahl, Burri, & Edwards, 2019). This lack of resolution severely challenges studies of linkage and recombination as well as inference of selection, and it is an Achilles heel of many contemporary studies of MHC genes in evolutionary ecology (Gaigher et al., 2016; O’Connor et al., 2019). The use of family data to infer segregating MHC haplotypes offers a powerful method to overcome these challenges (Gaigher et al., 2016, 2018). However, for such studies to capture a significant part of the MHC haplotype variation in wild populations, segregation patterns should be analyzed in a large number of families - especially when studying species with high levels of MHC diversity, such as songbirds (clade Passeri of Passeriformes) (Minias, Pikus, Whittingham, & Dunn, 2018; O’Connor, Strandh, Hasselquist, Nilsson, & Westerdahl, 2016). Fortunately, as the cost of high-throughput DNA sequencing continues to decrease, genotyping the number of samples necessary to infer MHC haplotypes in a large number of families is becoming feasible to many research groups.

To assist such studies, we here present the R package MHCtools that contains a set of functions for automated MHC haplotype inference from family data (Roved, 2019). Besides MHC haplotype inference, MHCtools contains functions that facilitate the bioinformatical steps involved in filtering large amplicon sequencing data sets and downstream data analysis (Table 1). In the present paper, we demonstrate the use of MHCtools to assist filtering of MHC class I (MHC-I) amplicon sequencing data and to carry out automated MHC-I haplotype inference using data from a wild population of great reed warblers (*Acrocephalus arundinaceus*), a songbird with highly duplicated MHC genes and extensive gene copy number variation between individuals (Roved, Hansson, Tarka, Hasselquist, & Westerdahl, 2018; Westerdahl, Wittzell, & von Schantz, 1999; Westerdahl, Wittzell, von Schantz, & Bensch, 2004). We used the resulting haplotype data set to estimate the degree of MHC-I gene copy number variation in this species, and the degree of recombination in the chromosomal region containing the MHC-I genes. Finally, we investigated whether tight linkage among the MHC-I genes has favored evolution of haplotypes that combine highly divergent MHC-I alleles.

**Table 1.** Overview of the functions included in MHCtools v. 1.2.1.

|  |  |
| --- | --- |
| **CalcPdist** | Calculates nucleotide or amino acid p-distances (i.e. the proportion of variable positions) from pairwise sequence comparisons and mean p-distances for each sample in a DADA2 sequence table. |
| **CreateFas** | Creates a FASTA file with all the sequences in a DADA2 sequence table. |
| **CreateSamplesFas** | Creates a set of FASTA files with the sequences present in each sample in a DADA2 sequence table. |
| **HpltFind** | Automatically infers major histocompatibility complex (MHC) haplotypes from the genotypes of parents and offspring in families. The functions **GetHpltTable()** and **GetHpltStats()** are designed to evaluate the output files. |
| **PapaDiv** | Calculates the joint major histocompatibility complex (MHC) diversity in parent pairs, taking into account alleles that are shared between the parents. The joint diversity in parent pairs is useful for heritability analyses in non-model species, where one wants to estimate the heritability of MHC diversity. The number of unique alleles in offspring may not be directly derived from the parental genotypes if some alleles are shared between the parents. |
| **ReplMatch** | Automatically compares technical replicates in an amplicon sequencing data set and reports the proportion of mismatches. The functions **GetReplTable()** and **GetReplStats()** are designed to evaluate the output files. |

## Methods

### Data set

We used empirical data on 141 adult males, 131 adult females, and 287 chicks from our long term study population of great reed warblers breeding at lake Kvismaren (59°10’N, 15°25’E) in southern Central Sweden (Hansson et al., 2018; Hasselquist, Montras-Janer, Tarka, & Hansson, 2017; Roved et al., 2018). The adult individuals in our data set have been observed, examined, and ringed in the period 1984–2004, and the chicks constitute the 1998 and 1999 cohorts from the same population, with addition of one nest from each of the years 1992 and 1996 (cf. Bensch et al., 1998; Hasselquist, 1998; Tarka et al., 2014). All territorial males and breeding females in our study population were mist-netted and marked with aluminum rings and unique combinations of color rings (Bensch et al., 1998). We located > 95% of all nests before fledging of the offspring and registered all breeding attempts (Bensch et al., 1998). 8–10 days after hatching we ringed the nestlings with an aluminum ring and a year-specific color ring. The paternity and maternity of all chicks were verified by molecular methods (Hansson, Hasselquist, & Bensch, 2004; Hasselquist, Bensch, & von Schantz, 1995). Approximately 3% of the total number of offspring resulted from extra-pair mating (Hansson et al., 2004; Hasselquist et al., 1995; Hasselquist, Bensch, & von Schantz, 1996), and these were assigned to their genetic parents in all analyses.

Fieldwork and DNA sampling were approved by the Malmö/Lund Animal Ethics Committee and the Swedish Bird Ringing Centre.

### DNA sampling and sequencing

We collected 20–80 µl blood from each individual by puncturing the brachial or tarsus vein. The blood samples were suspended in 500 µl SET buffer (0.15 M NaCl, 0.05 M TRIS, 0.001 M EDTA), and kept frozen at – 20°C until DNA extraction. Isolation of genomic DNA was carried out by standard phenol/chloroform-isoamylalcohol extraction (Sambrook, Fritsch, & Maniatis, 1989). The purified DNA was stored in 1×TE buffer and frozen at –50 or –80°C.

We amplified a 262 bp region of MHC-I exon 3 from 559 great reed warblers using previously designed primers: HNalla-HN46 (O’Connor et al., 2016; Westerdahl et al., 2004). Approximately half of the amplicons (samples from 88 adult males, 100 adult females, and 145 chicks) were sequenced in a Roche 454 GS FLX (F. Hoffmann-La Roche AG, Basel, Switzerland) following the manufacturer’s instructions at the Department of Biology, Lund University. See Roved et al. (2018) for details on experimental setup, tags, and PCR amplification in the 454-sequencing experiment. The remaining amplicons were sequenced in two independent runs using 300 bp paired-end sequencing in an Illumina MiSeq (Illumina Inc., San Diego, CA, USA) following the manufacturer’s instructions at the Department of Biology, Lund University. Samples from 23 adult males, 32 adult females, and 150 chicks (including 11 replicates from the 454-sequencing experiment) were included in the first run and a smaller batch with samples from 30 adult males were added in a second run. In both runs, great reed warbler samples were multiplexed with other samples in libraries comprising 384 samples. Details on tags, PCR amplification, and library preparation for the Illumina MiSeq sequencing experiment are provided in the SI (Supplementary Methods).

### Filtering and screening of sequencing data

The 454 sequencing data were demultiplexed using the software jMHC (Stuglik, Radwan, & Babik, 2011) and filtered according to the filtering protocol described in Galan, Guivier, Caraux, Charbonnel, & Cosson (2010). Subsequently, the data were manually screened to remove low-quality and artificial sequences, and non-functional alleles. See (Roved et al., 2018) for further details on the results and bioinformatics protocol of the 454 sequencing experiment.

The Illumina MiSeq sequencing outputs were trimmed to remove adapters, primers, and tag sequences using the software Cutadapt version 1.14 (Martin, 2011). The trimmed sequences were then filtered using the R package DADA2 version 1.4.0 (Callahan et al., 2016) in R version 3.4.2 (R Core Team, 2017). We inspected the error rate plots for deviations from the error rates expected given the observed Q-values, and employed the integrated function removoBimeraDenovo in DADA2 to remove chimeras. The machine learning algorithm employed by DADA2 to estimate the error rates associated with a data set requires the setting of a prior of maximum expected error rates, and the final output from DADA2 depends on this prior setting (Roved et al., unpublished data). Different combinations of primers and/or study species may produce different amounts of sequencing errors; hence, prior settings may be unpredictable for new data sets. Therefore, each study should perform a careful evaluation of the accuracy obtained with a number of different prior settings. Such an evaluation can be performed by comparing technical replicates in the data set.

After preliminary filtering of the first Illumina data set we identified 52 samples that had identical genotypes to one or more other samples (predominantly full siblings) in the data set, and grouped these samples into 25 replicate sets. We then optimized the filterAndTrim settings in DADA2 by comparing the replicate sets in 29 filtering runs with different settings. For each of these runs, we employed the functions ReplMatch and GetReplStats in the R package MHCtools version 1.2.1 to obtain repeatability estimates for the optimization of the filterAndTrim settings in DADA2. We identified the optimal filterAndTrim settings by evaluating the repeatability obtained with each setting. The settings providing the highest repeatability were: maxN = 0, maxEE fw = 0.1, maxEE rv = 0.1, and truncQ = 20. DADA2 inferred 295 sequences in the first Illumina data set with these settings. We then used the function CreateFas in MHCtools to create a fasta file with all the filtered sequences, and inspected these manually in BioEdit version 7.2.5 (Hall, 1999) to remove non-functional variants (sequences containing stop codons or indels that induced shifts in the reading frame). The remaining sequences were filtered by their relative abundance within each amplicon (per amplicon frequency). We again optimized the filtering threshold using repeatability estimates obtained by comparing the replicate sets using ReplMatch and GetReplStats in MHCtools. The per amplicon frequency threshold that gave the highest repeatability was 0.014. After filtering and screening, our first Illumina data set contained 226 unique DNA alleles in 205 samples. 197 alleles were found in both the first Illumina and the 454 sequencing runs, while 29 alleles were only found in the first Illumina run and 132 alleles were only found in the 454 sequencing run. The number of reads per sample was normally distributed with a mean of 20,920, a minimum of 12,532, and a maximum of 32,781.

The data from the second Illumina run was filtered and inspected using the same procedure and settings as the first Illumina data set, resulting in an initial inference of 200 sequences which, after filtering by per amplicon frequency and removal of two samples with low read numbers (0 and 3,150 reads, respectively) resulted in 162 unique DNA alleles in 31 samples. 32 of these alleles were not observed in the previous Illumina or 454 data sets. The number of reads per sample in the second Illumina data set was normally distributed with a mean of 22,492, a minimum of 12,062, and a maximum of 38,521.

After filtering and screening, we merged the Illumina MiSeq and 454 sequencing data sets for downstream analyses. Two genotyped samples were excluded from further analyses due to labelling errors. All DNA sequences were blasted against the NCBI database (http://blast.ncbi.nlm.nih.gov/Blast.cgi) and novel alleles were named following the MHC standardized nomenclature (Klein et al., 1990).

### Haplotype inference

We employed the HpltFind function in MHCtools to infer MHC-I haplotypes in our data set. We initially used the genotyping data from 372 samples from 67 families that constituted the 1998 and 1999 cohorts from our study population (*i.e*., all families from those years, in total 78 parents and 282 chicks), one family from 1991 (2 parents and 5 chicks), and one from 1996 (2 parents and 3 chicks). Hereafter, these families will be referred to as ‘the 69 families’.

Among the 390 MHC-I alleles that we observed in our data set, the five most common ones were present in 98%, 97%, 87%, 83%, and 50% of the samples, respectively. Such common alleles are often present in bothparents in families, and it can be difficult to ascertain their presence in one or both haplotypes in each parent through analysis of segregation patterns. To obtain the highest possible resolution, we therefore investigated segregation patterns across multiple generations, using the detailed pedigree of our study population (Hansson, Åkesson, Slate, & Pemberton, 2005; Tarka et al., 2014). Among 26 parents from the 69 families, we were able to trace ancestry up to five generations back, while the remaining parents were immigrants, for which no pedigree data were available (SI, Pedigree table). We investigated allele segregation patterns in 50 ancestral families of the 26 parents from the 69 families, that we were able to trace in our pedigree (hereafter referred to as ‘the ancestral families’) (SI, Pedigree table). This data included the genotypes of in total 80 adult individuals, of which 18 were also observed as parents in the 1998 and 1999 cohorts.

We inferred haplotypes based on allele segregation patterns by the following steps (illustrated with a flow chart in Fig. 1). The haplotype inference process on our data set is summarized in Table 2.

**Table 2.** Summary of the haplotype inference process.

|  |  |
| --- | --- |
| **Steps 1–2** |  |
| Initial number of putative haplotypes | 225 |
| Initial mean proportion of unresolved alleles in putative haplotypes | 0.446 |
| **Steps 3–4** |  |
| Number of putative haplotypes identical by descend to other haplotypes | 165 |
| Number of unresolved alleles verified in putative haplotypes | 358 |
| Number of alleles added to putative haplotypes (failed initial assignment) | 155 |
| Number of alleles removed from putative haplotypes (invalid initial assignment) | 121 |
| **Step 5** |  |
| Number of sequencing errors called as alleles, removed from individual samples | 15 |
| *- proportion of the total number of allele assignments to genotypes* | *0.0019* |
| Number of null alleles added to individual samples | 430 |
| *- proportion of the total number of allele assignments to genotypes* | *0.051* |
| **Step 6** |  |
| Number of putative haplotypes estimated to be homologous to other haplotypes | 41 |
| Number of unresolved alleles verified in putative haplotypes | 128 |
| Number of alleles added to putative haplotypes (failed initial assignment) | 7 |
| Number of alleles removed from putative haplotypes (invalid initial assignment) | 33 |
| **Final number of putative haplotypes** | **107** |
| **Final mean proportion of unresolved alleles in putative haplotypes** | **0.255** |

**Fig. 1.**
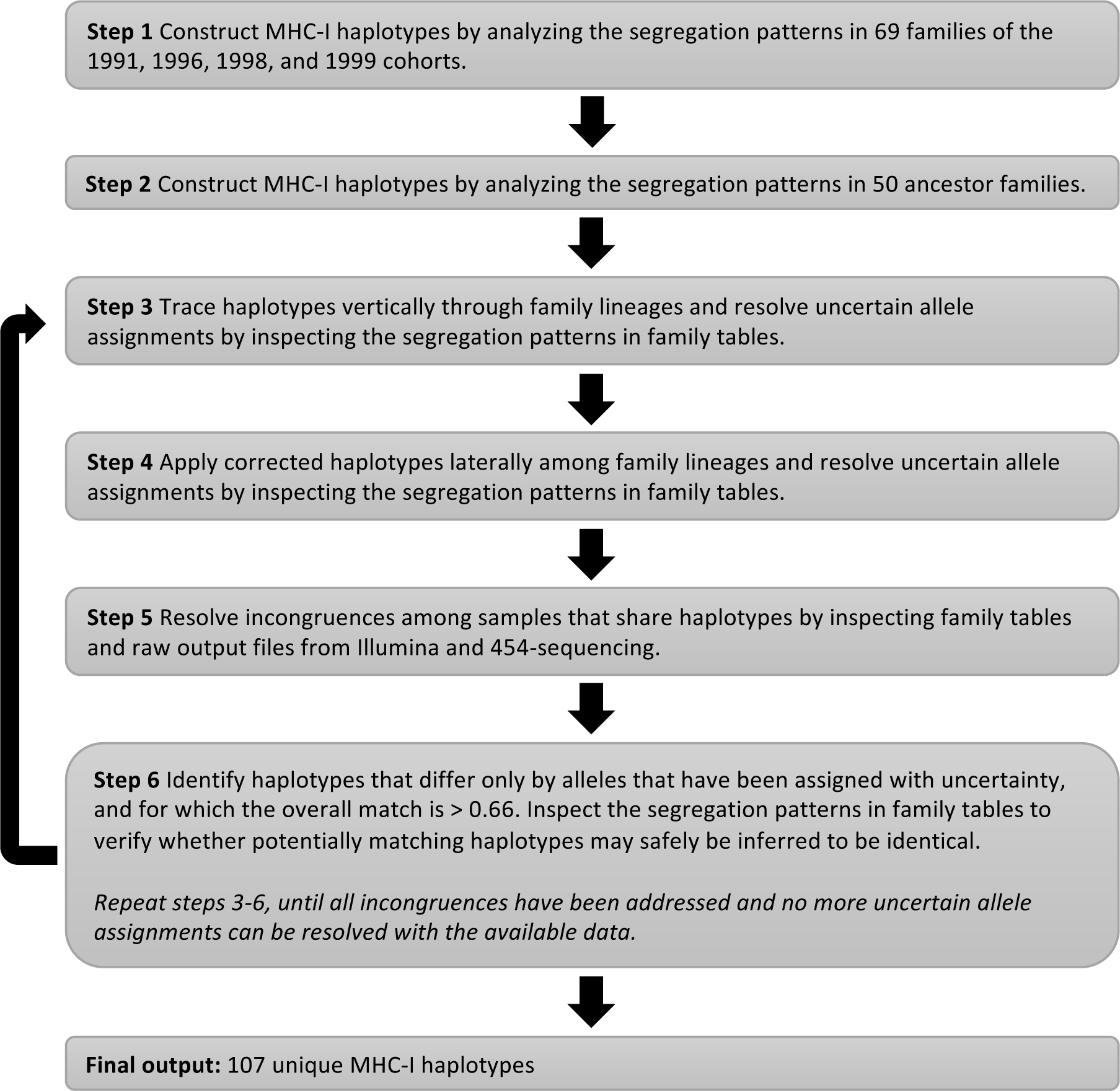
Flow chart of the haplotype inference protocol.

1. We used the HpltFind function in MHCtools to construct putative haplotypes for each individual in the 69 families. HpltFind assigns alleles to a parental haplotype if they occur in either parent and in one or more offspring. In cases when both parents and/or all or most offspring share an allele, it may not be possible to assign the allele to a parental haplotype with certainty. HpltFind assigns such alleles to all of their potential haplotypes and flags them as unresolved. We have illustrated this analysis for one family in Fig. 2.
2. We then constructed haplotypes for each individual in the ancestral families, using the same procedure as in step 1.
3. We traced the inheritance of all haplotypes through generations as far as data were available within each family lineage (SI, Pedigree table). When a haplotype mismatched between generations by alleles that had been flagged as unresolved, we inspected the family tables to verify whether we could resolve the allele assignments.
4. We then applied each corrected ancestral haplotype throughout all families in which it occurred, and resolved lateral mismatches between family lineages by the same procedure as we applied to the vertical mismatches observed within family lineages in step 3.
5. If incongruences were found between individuals that inherited or passed down a haplotype, and these incongruences could not be resolved by inspection of the family tables as described in steps 3–4, the most likely cause of the incongruence was identified. In rare cases, some alleles (i) had failed to amplify during PCR or had been erroneously deleted from samples in the filtering process (usually due to low PCR amplification success, *i.e*. null alleles), (ii) turned out to be sequencing errors that survived the filtering process (only alleles that had low read numbers and could be derived from more abundant alleles in the same sample by single nucleotide substitutions), or (iii) had not initially been assigned to a haplotype in the inference process (usually because it was a null allele in one or more samples in the family). Each incongruence was investigated by inspecting the family tables and the raw sequencing output files for presence or absence of the mismatching allele prior to filtering. In cases of multiple solutions to an incongruence, the solution involving the fewest assumptions was applied. By comparing haplotypes both within and between family lineages as described in steps 3–5, we were able to resolve a large proportion of the uncertain allele assignments (Table 2).
6. Finally, we identified potentially identical haplotypes throughout the data set, using a lower threshold of 0.66 for the proportion of matching sequences between any two haplotypes. We investigated all potential matches by manual inspection. Whenever a set of haplotypes only mismatched by alleles that had been assigned as uncertain (and had not been resolved in steps 3–5), we inspected the family tables to verify whether we could safely assume identity between the potentially matching haplotypes. This enabled us to resolve some additional uncertain allele assignments (Table 2).

**Fig. 2.**
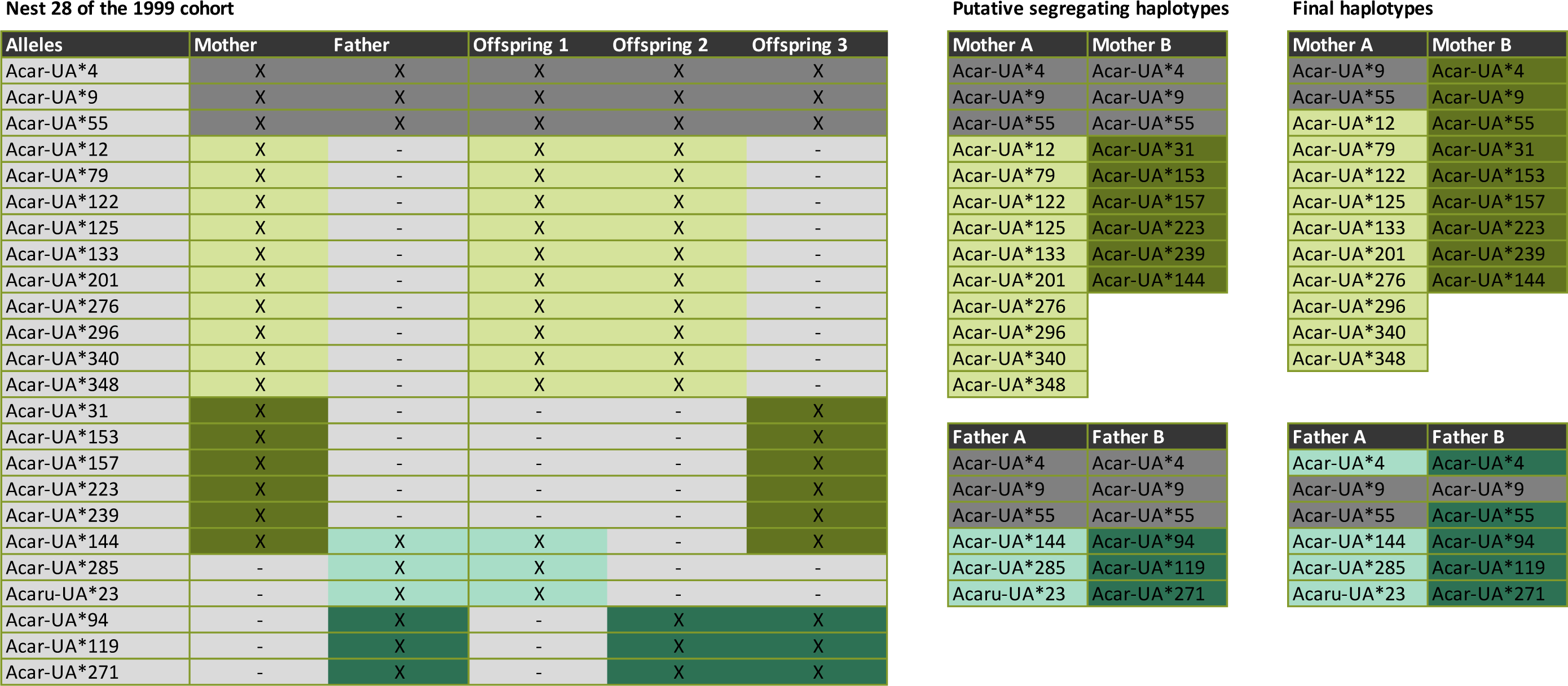
Family table from nest number 28 of the 1999 cohort showing MHC-I allele segregation patterns with inferred putative segregating MHC-I haplotypes (Mother A, Mother B, Father A, Father B) marked by different colors. Dark gray color indicates that a segregation pattern could not be determined for an allele, because it was present in both parents and in all offspring (uncertain allele). In the final haplotypes, a number of uncertain alleles were resolved by applying steps 3-6 in the haplotype inference protocol (Fig. 1).

We repeated steps 3–6, further improving the exactness of our haplotype inference by reapplying the corrected haplotypes both within and between families and family lineages. This process was repeated until all incongruences had been addressed and no more uncertain allele assignments could be resolved with the available data.

We were unable to resolve the segregation of alleles in three of the ancestral families, as the genotypes did not overlap. The segregation patterns suggested that blood samples from two individuals in these families have been exchanged or mislabeled, so these samples and nests were excluded from further analyses.

### Estimating the recombination rate

We calculated the recombination rate as the number of recombinant haplotypes divided by the total number of gametes (*i.e*., the number of offspring in families for which we successfully inferred haplotypes multiplied by two).

### Genetic divergence between alleles in MHC-I haplotypes

We used the CalcPdist function in MHCtools to quantify the proportion of nucleotide and amino acid differences (p-distances) between all pairs of MHC-I alleles, and calculated the means of the pairwise p-distances between all alleles in each haplotype.

To test whether alleles were more divergent within haplotypes than expected by chance, we generated 1,000 *in silico* simulations of our haplotype data set. The simulated data sets were generated by randomly assigning existing alleles to haplotypes while maintaining the number of different alleles for each haplotype. We calculated means of the pairwise nucleotide and amino acid p-distances between all alleles in each simulated haplotype and, for each simulation, calculated a mean across all haplotypes. Finally, for both nucleotide and amino acid p-distances, we compared the mean of the p-distances observed across all real haplotypes against the distribution of the mean p-distances derived from the simulated haplotype sets. The p-distance calculations, data simulations, and t-tests were carried out in R version 3.6.1 (R Core Team, 2019).

## Results

We developed a stepwise protocol for inference of MHC haplotypes in non-model species, based on the function HpltFind in the R package MHCtools (Roved, 2019), which carries out automated analysis of allele segregation patterns in family data (Fig. 1). We demonstrated the utility of this and a number of auxiliary functions from MHCtools by carrying out MHC-I genotyping and haplotype inference on an empirical data set of 559 great reed warblers from our long-term study population in Lake Kvismaren in southern Central Sweden.

We genotyped the individuals in our data set by amplifying and sequencing the MHC-I exon 3 using high-throughput Roche 454 and Illumina MiSeq platforms. The data set that was sequenced using the Roche 454 platform was filtered according to the principles in Galan et al. (2010), and we achieved a repeatability of 0.94 for these samples (estimated across 50 replicate sets) (Roved et al., 2018). The data set that was sequenced using Illumina MiSeq was filtered with DADA2 (Callahan et al., 2016), and we achieved a near perfect repeatability of 0.998 (estimated across 25 replicate sets). When comparing 11 replicated MHC-I genotypes between the Roche 454 and Illumina MiSeq runs, we achieved a repeatability of 0.96 between the sequencing platforms. The final, merged data set contained 390 unique MHC-I exon 3 alleles in 559 samples, of which 324 alleles were unique at the amino acid sequence level.

We successfully identified 107 unique MHC-I haplotypes based on allele segregation patterns in 116 great reed warbler families (SI, Haplotype Tables). Segregation patterns may be hard to resolve for alleles that are observed at high frequencies in the population, and our data set contained five MHC-I alleles with allele frequencies > 0.5. However, by analyzing allele segregation across multiple generations, we were able to reduce the mean proportion of unresolved alleles to 0.199 for haplotypes that were observed in multiple families, while it was 0.327 for haplotypes that were only observed in single families. For all samples, the mean proportion of unresolved alleles was 0.255 (Table 2). Furthermore, our haplotype inference process revealed a high congruence between the genotypes of related individuals in our data set. In the haplotype inference process, we discovered and removed 15 sequencing errors from individual samples in our data set (type I errors), corresponding to a proportion of 0.0019 of the total number of allele assignments (Table 2). We discovered 430 null alleles (type II errors), corresponding to a proportion of 0.051 of the total number of allele assignments, which were subsequently added to individual samples (Table 2). The larger number of type II compared to type I errors was expected, since the amplification of individual loci may be more or less successful in different samples.

We found considerable variation in the number of MHC-I gene copies between haplotypes, with as little as four and as many as 21 different alleles in single haplotypes. The mean number of different MHC-I alleles per haplotype was 9.2 with a standard deviation (S.D.) of 2.80 (Fig. 3). Furthermore, we found two recombinant haplotypes among the 334 offspring of the 116 families. From this observation, we estimated a recombination rate of 0.0030 within the genomic region spanned by the MHC-I in great reed warblers, corresponding to a distance of 0.3 centimorgan (cM).

**Fig. 3.**
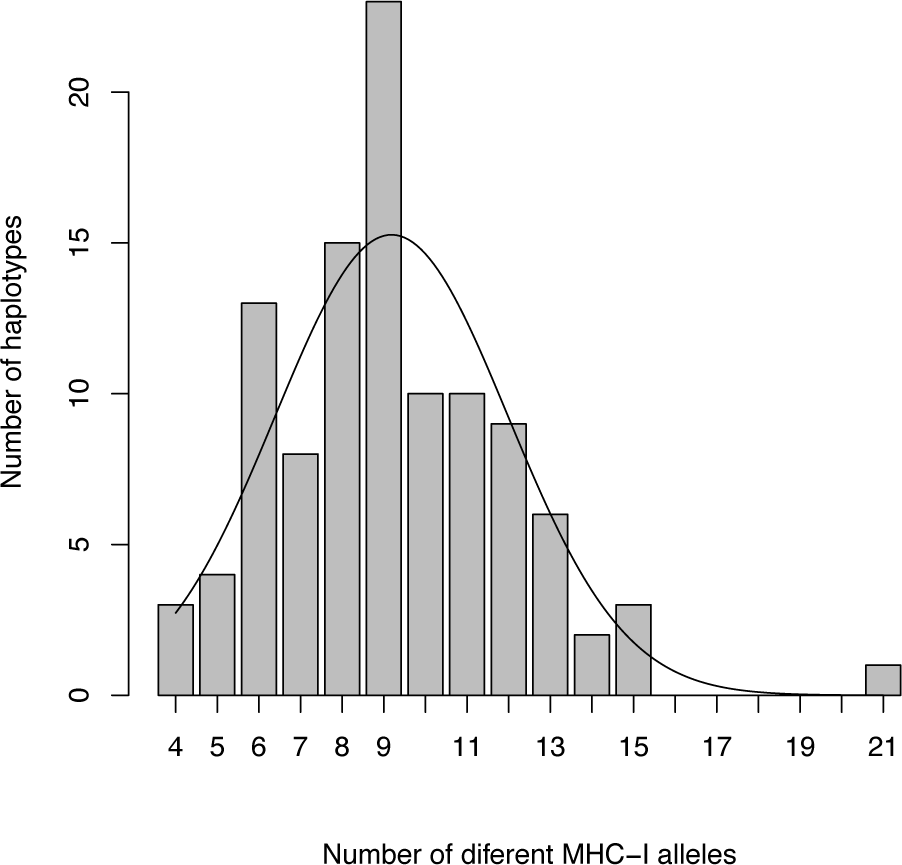
Distribution of the number of different MHC-I alleles on haplotypes. The line shows a normal distribution with the observed mean (9.2) and standard deviation (2.80).

The mean nucleotide divergence in the MHC-I haplotypes in our data set ranged from 0.083 to 0.139 with a mean of 0.118 (S.D. = 0.0097). The mean amino acid divergence in the MHC-I haplotypes in our data set ranged from 0.145 to 0.217 with a mean of 0.186 (S.D. = 0.0130). To investigate whether natural selection may have favored combinations of MHC-I alleles that are highly divergent from each other, we tested the mean nucleotide and amino acid p-distances observed in real haplotypes against similar measures obtained from 1,000 simulated data sets, in which alleles were randomly assigned to haplotypes. The mean p-distances across the real haplotypes were significantly larger than across the simulated haplotypes, both on the nucleotide and amino acid levels (p < 0.001 in both tests; Table 3; Fig. 4).

**Table 3.**
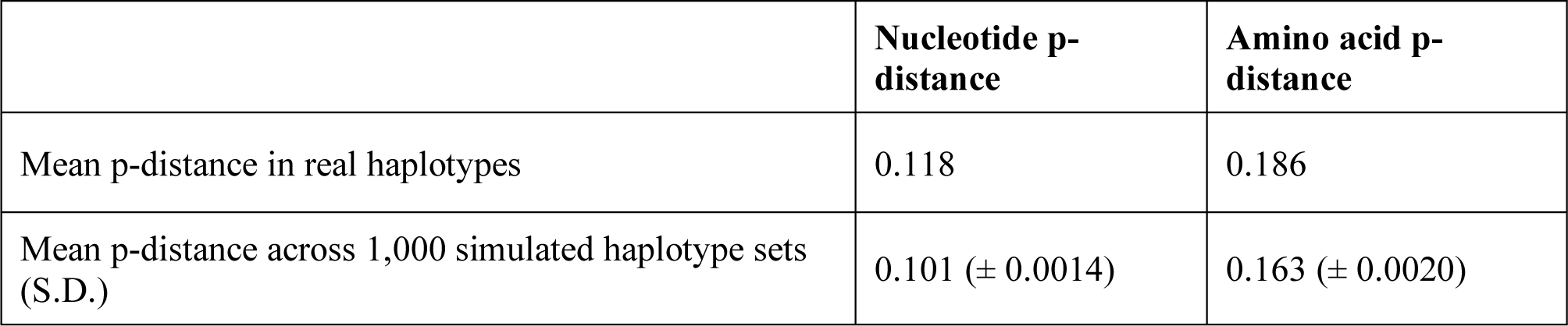
Observed mean nucleotide and amino acid p-distances in real haplotypes and across 1,000 simulated haplotype data sets.

|  | Nucleotide *p*-distance | Amino acid *p*-distance |
| --- | --- | --- |
| Mean *p*-distance in real haplotypes | 0.118 | 0.186 |
| Mean *p*-distance across 1,000 simulated haplotype sets (S.D.) | 0.101 ($\pm 0.0014$) | 0.163 ($\pm 0.0020$) |

**Fig. 4.**
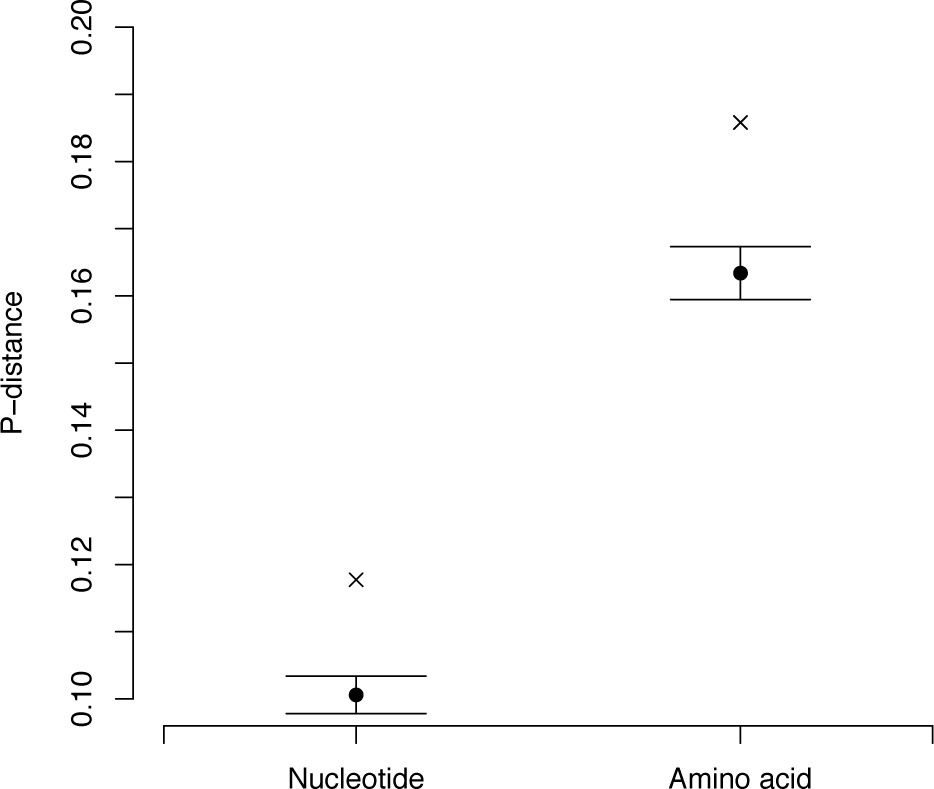
Observed mean nucleotide and amino acid p-distances on real haplotypes (crosses) versus mean nucleotide and amino acid p-distances ± 1 S.D. across 1,000 simulated haplotype data sets (dots w. error bars).

## Discussion

In the present study, we demonstrated the use of the newly developed R package MHCtools (Roved, 2019) for analysis of allele segregation patterns in species with highly duplicated gene families. We employed MHCtools in a stepwise protocol to characterize segregating MHC-I haplotypes in 116 families of a wild songbird, the great reed warbler. We identified 107 unique MHC-I haplotypes, and overall, observed a high degree of congruence in our haplotype inference process, which confirmed the accuracy of our genotyping methods. This provides a solid validation of our protocols, in particular given that haplotype inference is difficult in species with highly duplicated MHC genes (such as the great reed warbler) (Gaigher et al., 2018). The number of different MHC-I haplotypes that we observed in our population corresponds to the number observed in a previous study on barn owls, where 111 MHC-I haplotypes were observed among 140 families (Gaigher et al., 2018). Such standing genetic variation in MHC diversity could serve an important evolutionary function by enabling rapid adaptive shifts in response to the dynamics of faster evolving pathogens (Alves et al., 2019; O’Connor et al., 2019). Our analyses of the allele segregation patterns confirmed strong linkage of the MHC-I loci in the great reed warbler with a recombination rate of 0.0030, corresponding to a genetic distance of 0.3 cM.

As tightly linked MHC loci often co-segregate, investigations of the structure of MHC genes, haplotypes, and recombination between MHC loci may improve our understanding of correlations between MHC variation, fitness, and disease in wild populations (O’Connor et al., 2019). The MHC exhibits extraordinary evolutionary dynamics with rapid expansions and contractions of MHC gene copy number, and substantial variation in MHC sequence and haplotype structure (Kelley, Walter, & Trowsdale, 2005; Minias et al., 2018; Masatoshi Nei & Rooney, 2005; O’Connor et al., 2016; Ohta, 1991; Spurgin et al., 2011). Thus, previous studies have reported considerable variation in the number of different MHC alleles between individuals within species, suggesting that MHC gene copy number variation may be a common trait, at least among birds (Biedrzycka, O’Connor, et al., 2017; Gaigher et al., 2016; Roved et al., 2018; Stervander, Dierickx, Thorley, Brooke, & Westerdahl, 2020; Whittingham, Dunn, Freeman-Gallant, Taff, & Johnson, 2018). Our analyses of MHC-I haplotypes confirmed previous indications of substantial MHC-I gene copy number variation in the great reed warbler (O’Connor et al., 2016; Roved et al., 2018), with a minimum of 4 and a maximum of 21 different MHC-I alleles per haplotype (Fig. 3). Such variation in the number of different alleles on MHC haplotypes may seem puzzling, as one would expect selection to purge haplotypes that deviate too much from the optimal level of MHC diversity in the population. One explanation could be that variation in the number of alleles per haplotype is offset by variation either in the functional divergence between alleles or in the degree of specificity in the peptide binding properties of the alleles (cf. Chappell et al., 2015). It is also possible that deviations from optimal MHC diversity are offset by selective advantages associated with particular alleles (e.g. Bateson et al., 2016; Bonneaud, Perez-Tris, Federici, Chastel, & Sorci, 2006; Sepil, Lachish, Hinks, & Sheldon, 2013; Westerdahl et al., 2005), or that haplotypes with lower than optimal diversity are maintained in combinations with other haplotypes that harbor high diversity - and vice versa. Gene expression could also play a role, since not all MHC-I genes may be expressed to the same degree (Chappell et al., 2015; Drews, Strandh, Råberg, & Westerdahl, 2017). Furthermore, we have previously found evidence for a sexual conflict over MHC-I diversity in the great reed warbler, as having a higher than average number of different MHC-I alleles was advantageous for males, while the opposite was true for females (Roved et al., 2018). It is plausible that the selective advantage of having high MHC-I diversity in males could help maintain haplotypes with high numbers of alleles in great reed warblers, while, on the other hand, the disadvantage of such haplotypes in females could help maintain haplotypes with only few alleles.

Besides investigating MHC diversity in terms of the number of different alleles, we also took advantage of our haplotype data to investigate whether selection may have favored haplotypes that combine highly divergent MHC-I alleles in the great reed warbler. We showed that the mean proportions of nucleotide and amino acid differences between alleles were significantly higher in real haplotypes than in simulated haplotypes, to which alleles were assigned at random (Fig. 4). Because of the strong linkage among the MHC-I loci in the great reed warbler, non-random association of MHC-I alleles in haplotypes may be adaptive because it ensures that optimal MHC diversity is transmitted from parents to offspring. Our results suggest that selection favors combinations of highly divergent alleles in MHC-I haplotypes, because it is likely to increase the range of pathogens that can be recognized by the adaptive immune system, following the principle of the divergent allele advantage hypothesis (Wakeland et al., 1990). However, optimal individual MHC diversity in offspring may also be achieved by parents commonly choosing to mate with MHC-compatible partners (Aeschlimann, Haberli, Reusch, Boehm, & Milinski, 2003; Penn & Potts, 1999; Strandh et al., 2012). Even so, selection may favor non-random association of MHC alleles in haplotypes, because it may be advantageous for offspring irrespective of the degree of compatibility between maternally and paternally transmitted alleles. This may in particular be true under conditions that preclude random mating, e.g. in inbred or bottlenecked populations. Non-random association of highly divergent MHC alleles in haplotypes has previously been shown in wild chacma baboons (*Papio ursinus*), where it was suggested that selection favors haplotypes that combine MHC-DRB alleles with dissimilar physicochemical properties across multiple loci (Huchard et al., 2008). In contrast, Gaigher et al. (2018) found no evidence for a shift towards highly divergent allele combinations in MHC class I or II haplotypes in barn owls.

To our knowledge, this is the first study attempting to characterize MHC haplotypes in a species with highly duplicated MHC genes. However, while our results provide clear evidence for non-random association of MHC-I alleles in haplotypes in the great reed warbler, further investigations are required to establish whether alleles at different loci share the same evolutionary history. In humans, the MHC-I contains three highly polymorphic classical loci involved in antigen presentation (HLA-A, -B, and -C), and these have evolved divergent antigen binding properties (Buhler et al., 2016; Nei, Gu, & Sitnikova, 1997; Paul et al., 2013; Pierini & Lenz, 2018; Rao, De Boer, van Baarle, Maiers, & Kesmir, 2013). In birds, extraordinary divergence has been shown for MHC class IIB, where alleles separate into two lineages that predate the radiation of extant species (Goebel et al., 2017), but no such pattern has been described for MHC-I. To achieve a better understanding of the organization of and the evolutionary relationship between MHC-I genes in birds and other non-model species, it is necessary to investigate how different alleles associate with different loci. Understanding how MHC alleles segregate in haplotypes is an obvious step along that road, and may be complemented by analyses of the phylogenetic relationship between alleles, as exemplified in e.g. Goebel et al. (2017) and Burri et al. (2008), and by studies that attempt to map the physical structure of the MHC region (O’Connor et al., 2019). The latter may be possible to achieve by genome sequencing, in particular using long-read or linked read technologies, but current genome assembly methods have yet to overcome obstacles presented by the large number of duplicated sequences in the MHC region (Näpflin et al., 2019; O’Connor et al., 2019). Until such methods are developed and can be applied across a range of both species and individuals, linkage maps derived from haplotype inference may offer valuable insights into the structural organization of the MHC region. In addition, recent advances in studies on humans have demonstrated that incorporating prior knowledge about population-wide haplotype diversity may improve the accuracy of MHC genotyping in *de novo* genome assemblies (Dilthey, Cox, Iqbal, Nelson, & McVean, 2015). This underlines both the current and future value of methods and tools, such as those presented in this paper, that facilitate population-wide screening of MHC haplotypes in non-model species, where no such data are available *a priori*.

## Acknowledgements

We wish to thank A. Drews and T. Johansson for library constructions and high-troughput sequencing. Furthermore, we are grateful to S. Bensch, B. Nielsen, M. Åkesson, and staff at Kvismare Bird Observatory for considerable help with the great reed warbler fieldwork. Funding and support were provided by the Swedish Research Council (621-2013-4357, 2016-04391 to DH; 621-2014-5222, 2016-00689 to BH; 2015-05149 to HW), the European Research Council (ERC) under the European Union’s Horizon 2020 research and innovation program (Start Up grant 679799 to HW; Advanced grant 742646 to DH), a Linnaeus research excellence grant (349-2007-8690) from the Swedish Research Council and Lund University to the Centre for Animal Movement Research (CAnMove), Lunds Djurskyddsfond, Kungl. Fysiografiska Sällskapet i Lund (JR), and Kvismare Bird Observatory (report number 193).

## Data Accessibility

MHCtools version 1.2.1 (including user manual and documentation) is available at CRAN: https://cran.r-project.org/package=MHCtools. Our data set is available at the Zenodo repository: http://dx.doi.org/10.5281/zenodo.3716048 (Roved, Hansson, Stervander, Hasselquist, & Westerdahl, 2020). DNA sequences are available at GenBank: https://ncbi.nlm.nih.gov (accession numbers: MH468831–MH469159; MT193762– MT193822).

## Author contributions

JR designed the data analysis protocols and conceived, designed, and created the R package MHCtools; JR, BH, DH, and HW jointly conceived the study of MHC-I haplotypes in great reed warblers; MS constructed the amplicon sequencing libraries for Illumina sequencing; JR carried out bioinformatics, analyzed the data, and wrote the paper, with input from all authors.

