## Supplementary Information for "Non-random association of MHC-I alleles in favor of high diversity haplotypes in wild songbirds revealed by computer-assisted MHC haplotype inference using the R package MHCtools"

**Supplementary methods**

*Tags, PCR amplification, and library preparation for the Illumina MiSeq sequencing runs*
We prepared both amplicon libraries for Illumina sequencing in a two-step amplification. First, individual samples were amplified using the HNalla and HN46, modified with 5'-overhangs designed to match the Illumina sequencing adapters and molecular identifiers (MIDs) of the Nextera® XT v2 Index Kit (Illumina Inc., San Diego, CA, USA). The reactions comprised 25 µl and used 25 ng template DNA, 0.5 µM of each primer, and 12.5 µl 2X Phusion High-Fidelity PCR Master Mix (ThermoFisher Scientific, Waltham, USA). The PCR was initiated with a 30 s denaturation step at 98°C followed by 25 cycles of 10 s denaturation at 98°C, 10 s annealing at 66.8°C, and 15 s elongation at 72°C. A 10 min final extension 72°C completed the program.

The PCR product was cleaned with Agencourt AMPure XP-PCR Purification Kit (Beckman Coulter, Indianapolis, USA), following the manufacturer's instruction with some modifications: The ratio of PCR product to beads was 1:0.8, 80% ethanol was used in the bead cleaning steps, and the elution was made with 43 µl double-distilled water, which incubated at room temperature for two minutes. An aliquot of the clean PCR product was run on a 2% agarose gel, to verify fragment length and to roughly estimate concentration of the PCR product based on band intensity. The individual PCR products were then differentially evaporated at room temperature, to achieve even concentrations.

To be able to assign sequences to individual samples after multiplexing, we added unique combinations of forward and reverse Illumina indices to each sample using the Nextera XT v2 Index Kit (Illumina Inc., San Diego, CA, USA). A second PCR was run in 50 µl reactions that contained 25 µl 2X Phusion High-Fidelity PCR Master Mix (ThermoFisher Scientific, Waltham, USA), 5 µl of each index primer, and a varying amount of cleaned PCR product depending on estimated concentration (5, 10, or 15 µl for the first library; 5, 7.5, or 10 µl for the second library). The PCRs were initiated with a 30 s heating phase at 98°C, followed by eight cycles of 10 s denaturation at 98°C, 15 s annealing at 62°C, and 15 s elongation at 72°C, and ended with 10 min final extension at 72°C.

The indexed amplicons were cleaned with Agencourt AMPure XP-PCR Purification Kit (Beckman Coulter, Indianapolis, USA), following the manufacturer's instruction with some modifications: The ratio of PCR product to beads was 1:1.12, 80% ethanol was used in the bead cleaning steps, and the elution was made with 43 µl double-distilled water and incubated at room temperature for two minutes. The cleaned PCR products were checked on a 2% agarose gel, and quantified using a Quant-iT PicoGreen dsDNA Assay Kit (ThermoFisher Scientific/Invitrogen, Waltham, USA) modified for a 96-well plate, measured on a plate reader.

For each library, we pooled an equimolar quantity of each of 384 samples (including samples unrelated to this study) into pools (nine for the first library and four for the second library), depending on amplicon length, concentration, and primer combination. These pools were then quantified with Qubit Broad Range and High Sensitivity kits (ThermoFisher Scientific, Waltham, USA), after which we ran them on a Bioanalyzer DNA 2100 chip for validation of quality and size. In a final step, equimolar quantities of all pools were combined in a 20 nM library.

**Haplotype tables**

Unique haplotypes in our data set with the proportion of unresolved allele assignments on each haplotype. The tables show the alleles found on each haplotype, with specification of allele assignment status (1 = unresolved allele assignment, 0 = definite allele assignment).

| **Acar-HPLT*01** | | Prop. unresolved alleles | 0.071429 |
|---|---|---|---|
| Allele | Unresolved | | |
| Acar-UA*4 | 0 | | |
| Acar-UA*8 | 0 | | |
| Acar-UA*9 | 1 | | |
| Acar-UA*12 | 0 | | |
| Acar-UA*33 | 0 | | |
| Acar-UA*55 | 0 | | |
| Acar-UA*74 | 0 | | |
| Acar-UA*95 | 0 | | |
| Acar-UA*122 | 0 | | |
| Acar-UA*144 | 0 | | |
| Acar-UA*145 | 0 | | |
| Acar-UA*163 | 0 | | |
| Acar-UA*192 | 0 | | |
| Acar-UA*391 | 0 | | |

| **Acar-HPLT*02** | | Prop. unresolved alleles | 0.111111 |
|---|---|---|---|
| Allele | Unresolved | | |
| Acar-UA*9 | 1 | | |
| Acar-UA*48 | 0 | | |
| Acar-UA*55 | 0 | | |
| Acar-UA*82 | 0 | | |
| Acar-UA*126 | 0 | | |
| Acar-UA*145 | 0 | | |
| Acar-UA*157 | 0 | | |
| Acar-UA*241 | 0 | | |
| Acar-UA*330 | 0 | | |

| **Acar-HPLT*03** | | Prop. unresolved alleles | 0.1 |
|---|---|---|---|
| Allele | Unresolved | | |
| Acar-UA*9 | 1 | | |
| Acar-UA*54 | 0 | | |
| Acar-UA*130 | 0 | | |
| Acar-UA*133 | 0 | | |
| Acar-UA*153 | 0 | | |
| Acar-UA*157 | 0 | | |
| Acar-UA*201 | 0 | | |
| Acar-UA*223 | 0 | | |
| Acar-UA*239 | 0 | | |
| Acar-UA*296 | 0 | | |

**Acar-HPLT*04** Prop. unresolved alleles 0.142857

| Allele | Unresolved |
|---|---|
| Acar-UA*4 | 0 |
| Acar-UA*9 | 0 |
| Acar-UA*11 | 0 |
| Acar-UA*55 | 1 |
| Acar-UA*69 | 0 |
| Acar-UA*157 | 0 |
| Acar-UA*247 | 0 |

**Acar-HPLT*05** Prop. unresolved alleles 0

| Allele | Unresolved |
|---|---|
| Acar-UA*4 | 0 |
| Acar-UA*9 | 0 |
| Acar-UA*55 | 0 |
| Acar-UA*131 | 0 |
| Acar-UA*134 | 0 |
| Acar-UA*375 | 0 |

**Acar-HPLT*06** Prop. unresolved alleles 0.2

| Allele | Unresolved |
|---|---|
| Acar-UA*4 | 1 |
| Acar-UA*9 | 1 |
| Acar-UA*12 | 0 |
| Acar-UA*49 | 0 |
| Acar-UA*55 | 1 |
| Acar-UA*75 | 0 |
| Acar-UA*82 | 0 |
| Acar-UA*89 | 0 |
| Acar-UA*122 | 0 |
| Acar-UA*128 | 0 |
| Acar-UA*146 | 0 |
| Acar-UA*168 | 0 |
| Acar-UA*229 | 0 |
| Acar-UA*245 | 0 |
| Acar-UA*420 | 0 |

**Acar-HPLT*07** Prop. unresolved alleles 0.125

| Allele | Unresolved |
|---|---|
| Acar-UA*4 | 0 |
| Acar-UA*9 | 1 |
| Acar-UA*12 | 0 |
| Acaru-UA*23 | 0 |
| Acar-UA*33 | 0 |
| Acar-UA*55 | 0 |
| Acar-UA*144 | 0 |
| Acar-UA*146 | 0 |

| **Acar-HPLT*08** | | Prop. unresolved alleles | 0.2 |
|---|---|---|---|
| Allele | Unresolved | | |
| Acar-UA*4 | 0 | | |
| Acar-UA*9 | 1 | | |
| Acaru-UA*23 | 0 | | |
| Acar-UA*55 | 0 | | |
| Acar-UA*285 | 0 | | |

| **Acar-HPLT*09** | | Prop. unresolved alleles | 0.285714 |
|---|---|---|---|
| Allele | Unresolved | | |
| Acar-UA*4 | 0 | | |
| Acar-UA*9 | 1 | | |
| Acar-UA*35 | 0 | | |
| Acar-UA*55 | 1 | | |
| Acar-UA*66 | 0 | | |
| Acar-UA*192 | 0 | | |
| Acar-UA*265 | 0 | | |

| **Acar-HPLT*10** | | Prop. unresolved alleles | 0.4 |
|---|---|---|---|
| Allele | Unresolved | | |
| Acar-UA*4 | 0 | | |
| Acar-UA*9 | 1 | | |
| Acar-UA*31 | 0 | | |
| Acar-UA*47 | 0 | | |
| Acar-UA*55 | 1 | | |

| **Acar-HPLT*11** | | Prop. unresolved alleles | 0.25 |
|---|---|---|---|
| Allele | Unresolved | | |
| Acar-UA*9 | 1 | | |
| Acar-UA*31 | 0 | | |
| Acar-UA*55 | 1 | | |
| Acar-UA*78 | 0 | | |
| Acar-UA*208 | 0 | | |
| Acar-UA*265 | 0 | | |
| Acar-UA*357 | 0 | | |
| Acar-UA*397 | 0 | | |

| **Acar-HPLT*12** | | Prop. unresolved alleles | 0.076923 |
|---|---|---|---|
| Allele | Unresolved | | |
| Acar-UA*4 | 0 | | |
| Acar-UA*9 | 1 | | |
| Acar-UA*11 | 0 | | |
| Acar-UA*12 | 0 | | |
| Acar-UA*55 | 0 | | |
| Acar-UA*62 | 0 | | |
| Acar-UA*173 | 0 | | |
| Acar-UA*214 | 0 | | |
| Acar-UA*233 | 0 | | |
| Acar-UA*279 | 0 | | |
| Acar-UA*328 | 0 | | |
| Acar-UA*336 | 0 | | |
| Acar-UA*419 | 0 | | |

| **Acar-HPLT*13** | | Prop. unresolved alleles | 0.333333 |
|---|---|---|---|
| Allele | Unresolved | | |
| Acar-UA*4 | 0 | | |
| Acar-UA*9 | 1 | | |
| Acar-UA*31 | 0 | | |
| Acar-UA*36 | 0 | | |
| Acar-UA*55 | 1 | | |
| Acar-UA*144 | 0 | | |

| **Acar-HPLT*14** | | Prop. unresolved alleles | 0.111111 |
|---|---|---|---|
| Allele | Unresolved | | |
| Acar-UA*4 | 0 | | |
| Acar-UA*9 | 0 | | |
| Acar-UA*55 | 1 | | |
| Acar-UA*59 | 0 | | |
| Acar-UA*65 | 0 | | |
| Acar-UA*143 | 0 | | |
| Acar-UA*144 | 0 | | |
| Acar-UA*155 | 0 | | |
| Acar-UA*255 | 0 | | |

| **Acar-HPLT*15** | | Prop. unresolved alleles | 0.071429 |
|---|---|---|---|
| Allele | Unresolved | | |
| Acar-UA*9 | 1 | | |
| Acar-UA*51 | 0 | | |
| Acar-UA*52 | 0 | | |
| Acar-UA*55 | 0 | | |
| Acar-UA*107 | 0 | | |
| Acar-UA*110 | 0 | | |
| Acar-UA*143 | 0 | | |
| Acar-UA*144 | 0 | | |
| Acar-UA*186 | 0 | | |
| Acar-UA*231 | 0 | | |
| Acar-UA*255 | 0 | | |
| Acar-UA*304 | 0 | | |
| Acar-UA*368 | 0 | | |
| Acar-UA*392 | 0 | | |

| **Acar-HPLT*16** | | Prop. unresolved alleles | 0 |
|---|---|---|---|
| Allele | Unresolved | | |
| Acar-UA*9 | 0 | | |
| Acar-UA*37 | 0 | | |
| Acar-UA*55 | 0 | | |
| Acar-UA*60 | 0 | | |
| Acar-UA*94 | 0 | | |
| Acar-UA*144 | 0 | | |
| Acar-UA*224 | 0 | | |
| Acar-UA*250 | 0 | | |
| Acar-UA*349 | 0 | | |
| Acar-UA*350 | 0 | | |

| **Acar-HPLT*17** | | Prop. unresolved alleles | 0.285714 |
|---|---|---|---|
| Allele | Unresolved | | |
| Acar-UA*4 | 0 | | |
| Acar-UA*9 | 1 | | |
| Acar-UA*12 | 0 | | |
| Acar-UA*55 | 1 | | |
| Acar-UA*191 | 0 | | |
| Acar-UA*304 | 0 | | |
| Acar-UA*363 | 0 | | |

| **Acar-HPLT*18** | | Prop. unresolved alleles | 0.444444 |
|---|---|---|---|
| Allele | Unresolved | | |
| Acar-UA*9 | 1 | | |
| Acar-UA*12 | 0 | | |
| Acar-UA*31 | 1 | | |
| Acar-UA*48 | 0 | | |
| Acar-UA*55 | 1 | | |
| Acar-UA*86 | 0 | | |
| Acar-UA*122 | 0 | | |
| Acar-UA*144 | 1 | | |
| Acar-UA*330 | 0 | | |

| **Acar-HPLT*19** | | Prop. unresolved alleles | 0.166667 |
|---|---|---|---|
| Allele | Unresolved | | |
| Acar-UA*9 | 1 | | |
| Acar-UA*12 | 0 | | |
| Acar-UA*55 | 1 | | |
| Acar-UA*79 | 0 | | |
| Acar-UA*122 | 0 | | |
| Acar-UA*125 | 0 | | |
| Acar-UA*133 | 0 | | |
| Acar-UA*201 | 0 | | |
| Acar-UA*276 | 0 | | |
| Acar-UA*296 | 0 | | |
| Acar-UA*340 | 0 | | |
| Acar-UA*348 | 0 | | |

| **Acar-HPLT*20** | | Prop. unresolved alleles | 0.285714 |
|---|---|---|---|
| Allele | Unresolved | | |
| Acar-UA*4 | 1 | | |
| Acar-UA*9 | 1 | | |
| Acar-UA*30 | 0 | | |
| Acar-UA*39 | 0 | | |
| Acar-UA*54 | 0 | | |
| Acar-UA*61 | 0 | | |
| Acar-UA*144 | 0 | | |

| **Acar-HPLT*21** | | Prop. unresolved alleles | 0.125 |
|---|---|---|---|
| Allele | Unresolved | | |
| Acar-UA*4 | 0 | | |
| Acar-UA*9 | 1 | | |
| Acar-UA*12 | 0 | | |
| Acar-UA*136 | 0 | | |
| Acar-UA*144 | 0 | | |
| Acar-UA*191 | 0 | | |
| Acar-UA*241 | 0 | | |
| Acar-UA*364 | 0 | | |

| **Acar-HPLT*22** | | Prop. unresolved alleles | 0.25 |
|---|---|---|---|
| Allele | Unresolved | | |
| Acar-UA*4 | 0 | | |
| Acar-UA*9 | 1 | | |
| Acar-UA*12 | 0 | | |
| Acar-UA*55 | 1 | | |
| Acar-UA*77 | 0 | | |
| Acar-UA*144 | 0 | | |
| Acar-UA*223 | 0 | | |
| Acar-UA*238 | 0 | | |

| **Acar-HPLT*23** | | Prop. unresolved alleles | 0.333333 |
|---|---|---|---|
| Allele | Unresolved | | |
| Acar-UA*4 | 0 | | |
| Acar-UA*9 | 1 | | |
| Acar-UA*48 | 0 | | |
| Acar-UA*55 | 1 | | |
| Acar-UA*62 | 0 | | |
| Acar-UA*144 | 1 | | |
| Acar-UA*152 | 0 | | |
| Acar-UA*177 | 0 | | |
| Acar-UA*188 | 0 | | |

| **Acar-HPLT*24** | | Prop. unresolved alleles | 0.25 |
|---|---|---|---|
| Allele | Unresolved | | |
| Acar-UA*9 | 1 | | |
| Acar-UA*41 | 0 | | |
| Acar-UA*55 | 1 | | |
| Acar-UA*103 | 0 | | |
| Acar-UA*144 | 0 | | |
| Acar-UA*192 | 0 | | |
| Acar-UA*200 | 0 | | |
| Acar-UA*342 | 0 | | |

| **Acar-HPLT*25** | | Prop. unresolved alleles | 0.090909 |
|---|---|---|---|
| Allele | Unresolved | | |
| Acar-UA*4 | 0 | | |
| Acar-UA*8 | 0 | | |
| Acar-UA*9 | 0 | | |
| Acar-UA*12 | 0 | | |
| Acar-UA*48 | 0 | | |
| Acar-UA*55 | 1 | | |
| Acar-UA*58 | 0 | | |
| Acar-UA*144 | 0 | | |
| Acar-UA*145 | 0 | | |
| Acar-UA*236 | 0 | | |
| Acar-UA*304 | 0 | | |

| **Acar-HPLT*26** | | Prop. unresolved alleles | 0.125 |
|---|---|---|---|
| Allele | Unresolved | | |
| Acar-UA*9 | 1 | | |
| Acar-UA*12 | 0 | | |
| Acar-UA*47 | 0 | | |
| Acar-UA*137 | 0 | | |
| Acar-UA*144 | 0 | | |
| Acar-UA*208 | 0 | | |
| Acar-UA*279 | 0 | | |
| Acar-UA*280 | 0 | | |

| **Acar-HPLT*27** | | Prop. unresolved alleles | 0.111111 |
|---|---|---|---|
| Allele | Unresolved | | |
| Acar-UA*4 | 0 | | |
| Acar-UA*9 | 1 | | |
| Acar-UA*31 | 0 | | |
| Acar-UA*42 | 0 | | |
| Acar-UA*55 | 0 | | |
| Acar-UA*94 | 0 | | |
| Acar-UA*129 | 0 | | |
| Acar-UA*332 | 0 | | |
| Acar-UA*349 | 0 | | |

| **Acar-HPLT*28** | | Prop. unresolved alleles | 0.125 |
|---|---|---|---|
| Allele | Unresolved | | |
| Acar-UA*32 | 0 | | |
| Acar-UA*55 | 1 | | |
| Acar-UA*144 | 0 | | |
| Acar-UA*166 | 0 | | |
| Acar-UA*224 | 0 | | |
| Acar-UA*229 | 0 | | |
| Acar-UA*290 | 0 | | |
| Acar-UA*378 | 0 | | |

| **Acar-HPLT*29** | | Prop. unresolved alleles | 0 |
|---|---|---|---|
| Allele | Unresolved | | |
| Acar-UA*9 | 0 | | |
| Acar-UA*48 | 0 | | |
| Acar-UA*55 | 0 | | |
| Acar-UA*97 | 0 | | |
| Acar-UA*144 | 0 | | |
| Acar-UA*188 | 0 | | |
| Acar-UA*208 | 0 | | |
| Acar-UA*229 | 0 | | |
| Acar-UA*236 | 0 | | |
| Acar-UA*304 | 0 | | |

| **Acar-HPLT*30** | | Prop. unresolved alleles | 0 |
|---|---|---|---|
| Allele | Unresolved | | |
| Acar-UA*9 | 0 | | |
| Acar-UA*12 | 0 | | |
| Acar-UA*50 | 0 | | |
| Acar-UA*55 | 0 | | |
| Acar-UA*181 | 0 | | |
| Acar-UA*208 | 0 | | |
| Acar-UA*373 | 0 | | |

| **Acar-HPLT*31** | | Prop. unresolved alleles | 0.333333 |
|---|---|---|---|
| Allele | Unresolved | | |
| Acar-UA*4 | 0 | | |
| Acar-UA*9 | 1 | | |
| Acar-UA*55 | 1 | | |
| Acar-UA*123 | 0 | | |
| Acar-UA*144 | 1 | | |
| Acar-UA*149 | 0 | | |
| Acar-UA*183 | 0 | | |
| Acar-UA*196 | 0 | | |
| Acar-UA*401 | 0 | | |

| **Acar-HPLT*32** | | Prop. unresolved alleles | 0.363636 |
|---|---|---|---|
| Allele | Unresolved | | |
| Acar-UA*9 | 1 | | |
| Acar-UA*31 | 1 | | |
| Acar-UA*55 | 1 | | |
| Acar-UA*77 | 0 | | |
| Acar-UA*82 | 0 | | |
| Acar-UA*91 | 0 | | |
| Acar-UA*144 | 1 | | |
| Acar-UA*183 | 0 | | |
| Acar-UA*196 | 0 | | |
| Acar-UA*204 | 0 | | |
| Acar-UA*349 | 0 | | |

| **Acar-HPLT*33** | | Prop. unresolved alleles | 0.125 |
|---|---|---|---|
| Allele | Unresolved | | |
| Acar-UA*4 | 0 | | |
| Acar-UA*9 | 1 | | |
| Acar-UA*12 | 0 | | |
| Acar-UA*30 | 0 | | |
| Acar-UA*33 | 0 | | |
| Acar-UA*55 | 0 | | |
| Acar-UA*145 | 0 | | |
| Acar-UA*235 | 0 | | |

| **Acar-HPLT*34** | | Prop. unresolved alleles | 0.111111 |
|---|---|---|---|
| Allele | Unresolved | | |
| Acar-UA*4 | 0 | | |
| Acar-UA*9 | 1 | | |
| Acar-UA*43 | 0 | | |
| Acar-UA*48 | 0 | | |
| Acar-UA*55 | 0 | | |
| Acar-UA*190 | 0 | | |
| Acar-UA*236 | 0 | | |
| Acar-UA*304 | 0 | | |
| Acar-UA*367 | 0 | | |

| **Acar-HPLT*35** | | Prop. unresolved alleles | 0.272727 |
|---|---|---|---|
| Allele | Unresolved | | |
| Acar-UA*4 | 1 | | |
| Acar-UA*9 | 1 | | |
| Acar-UA*40 | 0 | | |
| Acar-UA*55 | 1 | | |
| Acar-UA*144 | 0 | | |
| Acar-UA*191 | 0 | | |
| Acar-UA*281 | 0 | | |
| Acar-UA*290 | 0 | | |
| Acar-UA*310 | 0 | | |
| Acar-UA*380 | 0 | | |
| Acar-UA*384 | 0 | | |

| **Acar-HPLT*36** | | Prop. unresolved alleles | 0.1 |
|---|---|---|---|
| Allele | Unresolved | | |
| Acar-UA*9 | 1 | | |
| Acar-UA*31 | 0 | | |
| Acar-UA*48 | 0 | | |
| Acar-UA*122 | 0 | | |
| Acar-UA*123 | 0 | | |
| Acar-UA*145 | 0 | | |
| Acar-UA*217 | 0 | | |
| Acar-UA*241 | 0 | | |
| Acar-UA*306 | 0 | | |
| Acar-UA*325 | 0 | | |

| **Acar-HPLT*37** | | Prop. unresolved alleles | 0.5 |
|---|---|---|---|
| Allele | Unresolved | | |
| Acar-UA*4 | 1 | | |
| Acar-UA*9 | 1 | | |
| Acar-UA*11 | 0 | | |
| Acar-UA*55 | 1 | | |
| Acar-UA*144 | 0 | | |
| Acar-UA*183 | 0 | | |

| **Acar-HPLT*38** | | Prop. unresolved alleles | 0.5 |
|---|---|---|---|
| Allele | Unresolved | | |
| Acar-UA*4 | 1 | | |
| Acar-UA*9 | 1 | | |
| Acar-UA*12 | 0 | | |
| Acar-UA*55 | 1 | | |
| Acar-UA*144 | 1 | | |
| Acar-UA*223 | 0 | | |
| Acar-UA*270 | 0 | | |
| Acar-UA*279 | 0 | | |

| **Acar-HPLT*39** | | Prop. unresolved alleles | 0.444444 |
|---|---|---|---|
| Allele | Unresolved | | |
| Acar-UA*4 | 1 | | |
| Acar-UA*8 | 0 | | |
| Acar-UA*9 | 1 | | |
| Acar-UA*55 | 1 | | |
| Acar-UA*73 | 0 | | |
| Acar-UA*144 | 1 | | |
| Acar-UA*156 | 0 | | |
| Acar-UA*175 | 0 | | |
| Acar-UA*225 | 0 | | |

| **Acar-HPLT*40** | | Prop. unresolved alleles | 0.333333 |
|---|---|---|---|
| Allele | Unresolved | | |
| Acar-UA*4 | 0 | | |
| Acar-UA*9 | 1 | | |
| Acaru-UA*23 | 0 | | |
| Acar-UA*31 | 0 | | |
| Acar-UA*55 | 1 | | |
| Acar-UA*144 | 1 | | |
| Acar-UA*146 | 0 | | |
| Acar-UA*277 | 0 | | |
| Acar-UA*322 | 0 | | |

| **Acar-HPLT*41** | | Prop. unresolved alleles | 0.25 |
|---|---|---|---|
| Allele | Unresolved | | |
| Acar-UA*4 | 0 | | |
| Acar-UA*9 | 1 | | |
| Acar-UA*12 | 0 | | |
| Acar-UA*55 | 1 | | |
| Acar-UA*122 | 0 | | |
| Acar-UA*190 | 0 | | |
| Acar-UA*192 | 0 | | |
| Acar-UA*235 | 0 | | |

| **Acar-HPLT*42** | | Prop. unresolved alleles | 0 |
|---|---|---|---|
| Allele | Unresolved | | |
| Acar-UA*137 | 0 | | |
| Acar-UA*144 | 0 | | |
| Acar-UA*187 | 0 | | |
| Acar-UA*208 | 0 | | |

| **Acar-HPLT*43** | | Prop. unresolved alleles | 0.090909 |
|---|---|---|---|
| Allele | Unresolved | | |
| Acar-UA*9 | 1 | | |
| Acar-UA*48 | 0 | | |
| Acar-UA*55 | 0 | | |
| Acar-UA*122 | 0 | | |
| Acar-UA*144 | 0 | | |
| Acar-UA*164 | 0 | | |
| Acar-UA*208 | 0 | | |
| Acar-UA*229 | 0 | | |
| Acar-UA*304 | 0 | | |
| Acar-UA*330 | 0 | | |
| Acar-UA*408 | 0 | | |

| **Acar-HPLT*44** | | Prop. unresolved alleles | 0.111111 |
|---|---|---|---|
| Allele | Unresolved | | |
| Acar-UA*4 | 0 | | |
| Acar-UA*9 | 1 | | |
| Acar-UA*55 | 0 | | |
| Acar-UA*128 | 0 | | |
| Acar-UA*144 | 0 | | |
| Acar-UA*191 | 0 | | |
| Acar-UA*337 | 0 | | |
| Acar-UA*371 | 0 | | |
| Acar-UA*410 | 0 | | |

| **Acar-HPLT*45** | | Prop. unresolved alleles | 0.166667 |
|---|---|---|---|
| Allele | Unresolved | | |
| Acar-UA*4 | 0 | | |
| Acar-UA*9 | 1 | | |
| Acar-UA*44 | 0 | | |
| Acar-UA*45 | 0 | | |
| Acar-UA*55 | 0 | | |
| Acar-UA*347 | 0 | | |

| **Acar-HPLT*46** | | Prop. unresolved alleles | 0.333333 |
|---|---|---|---|
| Allele | Unresolved | | |
| Acar-UA*4 | 1 | | |
| Acar-UA*9 | 1 | | |
| Acar-UA*31 | 0 | | |
| Acar-UA*55 | 1 | | |
| Acar-UA*73 | 0 | | |
| Acar-UA*114 | 0 | | |
| Acar-UA*115 | 0 | | |
| Acar-UA*144 | 0 | | |
| Acar-UA*159 | 0 | | |

| **Acar-HPLT*47** | | Prop. unresolved alleles | 0.1 |
|---|---|---|---|
| Allele | Unresolved | | |
| Acar-UA*4 | 0 | | |
| Acar-UA*9 | 0 | | |
| Acar-UA*12 | 0 | | |
| Acar-UA*55 | 1 | | |
| Acar-UA*144 | 0 | | |
| Acar-UA*208 | 0 | | |
| Acar-UA*279 | 0 | | |
| Acar-UA*280 | 0 | | |
| Acar-UA*318 | 0 | | |
| Acar-UA*326 | 0 | | |

| **Acar-HPLT*48** | | Prop. unresolved alleles | 0 |
|---|---|---|---|
| Allele | Unresolved | | |
| Acar-UA*4 | 0 | | |
| Acar-UA*9 | 0 | | |
| Acar-UA*31 | 0 | | |
| Acar-UA*55 | 0 | | |
| Acar-UA*144 | 0 | | |
| Acar-UA*153 | 0 | | |
| Acar-UA*157 | 0 | | |
| Acar-UA*223 | 0 | | |
| Acar-UA*239 | 0 | | |

| **Acar-HPLT*49** | | Prop. unresolved alleles | 0.333333 |
|---|---|---|---|
| Allele | Unresolved | | |
| Acar-UA*6 | 0 | | |
| Acar-UA*9 | 1 | | |
| Acar-UA*33 | 0 | | |
| Acar-UA*55 | 1 | | |
| Acar-UA*144 | 0 | | |
| Acar-UA*187 | 0 | | |

| **Acar-HPLT*50** | | Prop. unresolved alleles | 0.083333 |
|---|---|---|---|
| Allele | Unresolved | | |
| Acar-UA*9 | 1 | | |
| Acar-UA*12 | 0 | | |
| Acar-UA*41 | 0 | | |
| Acar-UA*55 | 0 | | |
| Acar-UA*107 | 0 | | |
| Acar-UA*143 | 0 | | |
| Acar-UA*144 | 0 | | |
| Acar-UA*186 | 0 | | |
| Acar-UA*231 | 0 | | |
| Acar-UA*255 | 0 | | |
| Acar-UA*277 | 0 | | |
| Acar-UA*368 | 0 | | |

| **Acar-HPLT*51** | | Prop. unresolved alleles | 0.181818 |
|---|---|---|---|
| Allele | Unresolved | | |
| Acar-UA*4 | 1 | | |
| Acar-UA*12 | 0 | | |
| Acar-UA*50 | 0 | | |
| Acar-UA*54 | 0 | | |
| Acar-UA*55 | 1 | | |
| Acar-UA*128 | 0 | | |
| Acar-UA*144 | 0 | | |
| Acar-UA*250 | 0 | | |
| Acar-UA*288 | 0 | | |
| Acar-UA*304 | 0 | | |
| Acar-UA*351 | 0 | | |

| **Acar-HPLT*52** | | Prop. unresolved alleles | 0.333333 |
|---|---|---|---|
| Allele | Unresolved | | |
| Acar-UA*4 | 1 | | |
| Acar-UA*9 | 1 | | |
| Acar-UA*49 | 0 | | |
| Acar-UA*55 | 1 | | |
| Acar-UA*81 | 0 | | |
| Acar-UA*122 | 0 | | |
| Acar-UA*128 | 0 | | |
| Acar-UA*144 | 1 | | |
| Acar-UA*147 | 0 | | |
| Acar-UA*160 | 0 | | |
| Acar-UA*187 | 0 | | |
| Acar-UA*229 | 0 | | |

| **Acar-HPLT*53** | | Prop. unresolved alleles | 0.166667 |
|---|---|---|---|
| Allele | Unresolved | | |
| Acar-UA*4 | 0 | | |
| Acar-UA*9 | 1 | | |
| Acar-UA*11 | 0 | | |
| Acar-UA*12 | 0 | | |
| Acar-UA*48 | 0 | | |
| Acar-UA*50 | 0 | | |
| Acar-UA*53 | 0 | | |
| Acar-UA*55 | 1 | | |
| Acar-UA*91 | 0 | | |
| Acar-UA*241 | 0 | | |
| Acar-UA*330 | 0 | | |
| Acar-UA*336 | 0 | | |

| **Acar-HPLT*54** | | Prop. unresolved alleles | 0.375 |
|---|---|---|---|
| Allele | Unresolved | | |
| Acar-UA*4 | 1 | | |
| Acar-UA*9 | 1 | | |
| Acar-UA*30 | 0 | | |
| Acar-UA*48 | 0 | | |
| Acar-UA*55 | 1 | | |
| Acar-UA*57 | 0 | | |
| Acar-UA*144 | 0 | | |
| Acar-UA*145 | 0 | | |

| **Acar-HPLT*55** | | Prop. unresolved alleles | 0.2 |
|---|---|---|---|
| Allele | Unresolved | | |
| Acar-UA*4 | 0 | | |
| Acar-UA*9 | 1 | | |
| Acar-UA*31 | 1 | | |
| Acar-UA*48 | 0 | | |
| Acar-UA*55 | 0 | | |
| Acar-UA*122 | 0 | | |
| Acar-UA*144 | 0 | | |
| Acar-UA*164 | 0 | | |
| Acar-UA*304 | 0 | | |
| Acar-UA*330 | 0 | | |

| **Acar-HPLT*56** | | Prop. unresolved alleles | 0.090909 |
|---|---|---|---|
| Allele | Unresolved | | |
| Acar-UA*4 | 0 | | |
| Acar-UA*12 | 0 | | |
| Acar-UA*48 | 0 | | |
| Acar-UA*50 | 0 | | |
| Acar-UA*54 | 0 | | |
| Acar-UA*128 | 0 | | |
| Acar-UA*144 | 1 | | |
| Acar-UA*250 | 0 | | |
| Acar-UA*288 | 0 | | |
| Acar-UA*304 | 0 | | |
| Acar-UA*351 | 0 | | |

| **Acar-HPLT*57** | | Prop. unresolved alleles | 0 |
|---|---|---|---|
| Allele | Unresolved | | |
| Acar-UA*4 | 0 | | |
| Acar-UA*9 | 0 | | |
| Acar-UA*55 | 0 | | |
| Acar-UA*144 | 0 | | |
| Acar-UA*158 | 0 | | |
| Acar-UA*368 | 0 | | |

| **Acar-HPLT*58** | | Prop. unresolved alleles | 0.230769 |
|---|---|---|---|
| Allele | Unresolved | | |
| Acar-UA*4 | 1 | | |
| Acar-UA*9 | 1 | | |
| Acar-UA*12 | 0 | | |
| Acar-UA*30 | 0 | | |
| Acar-UA*55 | 1 | | |
| Acar-UA*57 | 0 | | |
| Acar-UA*91 | 0 | | |
| Acar-UA*144 | 0 | | |
| Acar-UA*274 | 0 | | |
| Acar-UA*282 | 0 | | |
| Acar-UA*303 | 0 | | |
| Acar-UA*421 | 0 | | |
| Acar-UA*424 | 0 | | |

| **Acar-HPLT*59** | | Prop. unresolved alleles | 0.428571 |
|---|---|---|---|
| Allele | Unresolved | | |
| Acar-UA*4 | 1 | | |
| Acar-UA*9 | 1 | | |
| Acar-UA*12 | 0 | | |
| Acaru-UA*23 | 0 | | |
| Acar-UA*55 | 1 | | |
| Acar-UA*122 | 0 | | |
| Acar-UA*277 | 0 | | |

| **Acar-HPLT*60** | | Prop. unresolved alleles | 0.166667 |
|---|---|---|---|
| Allele | Unresolved | | |
| Acar-UA*4 | 0 | | |
| Acar-UA*9 | 1 | | |
| Acar-UA*55 | 0 | | |
| Acar-UA*94 | 0 | | |
| Acar-UA*119 | 0 | | |
| Acar-UA*271 | 0 | | |

| **Acar-HPLT*61** | | Prop. unresolved alleles | 0.2 |
|---|---|---|---|
| Allele | Unresolved | | |
| Acar-UA*4 | 0 | | |
| Acar-UA*9 | 1 | | |
| Acar-UA*55 | 0 | | |
| Acar-UA*144 | 0 | | |
| Acar-UA*200 | 0 | | |

| **Acar-HPLT*62** | | Prop. unresolved alleles | 0.375 |
|---|---|---|---|
| Allele | Unresolved | | |
| Acar-UA*4 | 0 | | |
| Acar-UA*9 | 1 | | |
| Acar-UA*50 | 0 | | |
| Acar-UA*53 | 0 | | |
| Acar-UA*55 | 1 | | |
| Acar-UA*144 | 1 | | |
| Acar-UA*184 | 0 | | |
| Acar-UA*308 | 0 | | |

| **Acar-HPLT*63** | | Prop. unresolved alleles | 0.2 |
|---|---|---|---|
| Allele | Unresolved | | |
| Acar-UA*4 | 0 | | |
| Acar-UA*9 | 1 | | |
| Acar-UA*12 | 0 | | |
| Acar-UA*30 | 0 | | |
| Acar-UA*31 | 0 | | |
| Acar-UA*55 | 1 | | |
| Acar-UA*122 | 0 | | |
| Acar-UA*144 | 1 | | |
| Acar-UA*145 | 0 | | |
| Acar-UA*157 | 0 | | |
| Acar-UA*208 | 0 | | |
| Acar-UA*240 | 0 | | |
| Acar-UA*278 | 0 | | |
| Acar-UA*315 | 0 | | |
| Acar-UA*327 | 0 | | |

| **Acar-HPLT*64** | | Prop. unresolved alleles | 0.333333 |
|---|---|---|---|
| Allele | Unresolved | | |
| Acar-UA*4 | 1 | | |
| Acar-UA*9 | 1 | | |
| Acar-UA*40 | 0 | | |
| Acar-UA*55 | 1 | | |
| Acar-UA*128 | 0 | | |
| Acar-UA*144 | 1 | | |
| Acar-UA*160 | 0 | | |
| Acar-UA*304 | 0 | | |
| Acar-UA*342 | 0 | | |
| Acar-UA*384 | 0 | | |
| Acar-UA*406 | 0 | | |
| Acar-UA*422 | 0 | | |

| **Acar-HPLT*65** | | Prop. unresolved alleles | 0.266667 |
|---|---|---|---|
| Allele | Unresolved | | |
| Acar-UA*4 | 1 | | |
| Acar-UA*9 | 1 | | |
| Acar-UA*12 | 0 | | |
| Acar-UA*48 | 0 | | |
| Acar-UA*55 | 1 | | |
| Acar-UA*72 | 0 | | |
| Acar-UA*91 | 0 | | |
| Acar-UA*122 | 0 | | |
| Acar-UA*144 | 1 | | |
| Acar-UA*145 | 0 | | |
| Acar-UA*148 | 0 | | |
| Acar-UA*268 | 0 | | |
| Acar-UA*304 | 0 | | |
| Acar-UA*336 | 0 | | |
| Acar-UA*384 | 0 | | |

| **Acar-HPLT*66** | | Prop. unresolved alleles | 0.333333 |
|---|---|---|---|
| Allele | Unresolved | | |
| Acar-UA*9 | 1 | | |
| Acar-UA*55 | 1 | | |
| Acar-UA*124 | 0 | | |
| Acar-UA*138 | 0 | | |
| Acar-UA*144 | 0 | | |
| Acar-UA*208 | 0 | | |

| **Acar-HPLT*67** | | Prop. unresolved alleles | 0.4 |
|---|---|---|---|
| Allele | Unresolved | | |
| Acar-UA*4 | 0 | | |
| Acar-UA*9 | 1 | | |
| Acar-UA*55 | 1 | | |
| Acar-UA*144 | 0 | | |
| Acar-UA*187 | 0 | | |

| **Acar-HPLT*68** | | Prop. unresolved alleles | 0.75 |
|---|---|---|---|
| Allele | Unresolved | | |
| Acar-UA*4 | 1 | | |
| Acar-UA*9 | 1 | | |
| Acar-UA*55 | 1 | | |
| Acar-UA*365 | 0 | | |

| **Acar-HPLT*69** | | Prop. unresolved alleles | 0.2 |
|---|---|---|---|
| Allele | Unresolved | | |
| Acar-UA*4 | 0 | | |
| Acar-UA*9 | 1 | | |
| Acar-UA*48 | 0 | | |
| Acar-UA*55 | 1 | | |
| Acar-UA*82 | 0 | | |
| Acar-UA*126 | 0 | | |
| Acar-UA*145 | 0 | | |
| Acar-UA*157 | 0 | | |
| Acar-UA*241 | 0 | | |
| Acar-UA*330 | 0 | | |

| **Acar-HPLT*70** | | Prop. unresolved alleles | 0.571429 |
|---|---|---|---|
| Allele | Unresolved | | |
| Acar-UA*4 | 0 | | |
| Acar-UA*9 | 1 | | |
| Acar-UA*31 | 0 | | |
| Acar-UA*55 | 1 | | |
| Acar-UA*144 | 1 | | |
| Acar-UA*260 | 0 | | |
| Acar-UA*304 | 1 | | |

| **Acar-HPLT*71** | | Prop. unresolved alleles | 0.307692 |
|---|---|---|---|
| Allele | Unresolved | | |
| Acar-UA*4 | 0 | | |
| Acar-UA*9 | 1 | | |
| Acar-UA*12 | 0 | | |
| Acar-UA*55 | 1 | | |
| Acar-UA*82 | 0 | | |
| Acar-UA*122 | 0 | | |
| Acar-UA*144 | 1 | | |
| Acar-UA*147 | 0 | | |
| Acar-UA*160 | 0 | | |
| Acar-UA*208 | 0 | | |
| Acar-UA*233 | 0 | | |
| Acar-UA*304 | 1 | | |
| Acar-UA*384 | 0 | | |

| **Acar-HPLT*72** | | Prop. unresolved alleles | 0.333333 |
|---|---|---|---|
| Allele | Unresolved | | |
| Acar-UA*4 | 1 | | |
| Acar-UA*9 | 1 | | |
| Acar-UA*51 | 0 | | |
| Acar-UA*107 | 0 | | |
| Acar-UA*136 | 0 | | |
| Acar-UA*143 | 0 | | |
| Acar-UA*144 | 1 | | |
| Acar-UA*304 | 0 | | |
| Acar-UA*368 | 0 | | |

| **Acar-HPLT*73** | | Prop. unresolved alleles | 0.272727 |
|---|---|---|---|
| Allele | Unresolved | | |
| Acar-UA*4 | 1 | | |
| Acar-UA*9 | 1 | | |
| Acar-UA*42 | 0 | | |
| Acar-UA*55 | 1 | | |
| Acar-UA*59 | 0 | | |
| Acar-UA*65 | 0 | | |
| Acar-UA*129 | 0 | | |
| Acar-UA*143 | 0 | | |
| Acar-UA*144 | 0 | | |
| Acar-UA*155 | 0 | | |
| Acar-UA*255 | 0 | | |

| **Acar-HPLT*74** | | Prop. unresolved alleles | 0.142857 |
|---|---|---|---|
| Allele | Unresolved | | |
| Acar-UA*4 | 0 | | |
| Acar-UA*9 | 0 | | |
| Acar-UA*55 | 0 | | |
| Acar-UA*144 | 1 | | |
| Acar-UA*199 | 0 | | |
| Acar-UA*254 | 0 | | |
| Acar-UA*394 | 0 | | |

| **Acar-HPLT*75** | | Prop. unresolved alleles | 0.111111 |
|---|---|---|---|
| Allele | Unresolved | | |
| Acar-UA*6 | 0 | | |
| Acar-UA*9 | 0 | | |
| Acar-UA*48 | 0 | | |
| Acar-UA*55 | 0 | | |
| Acar-UA*97 | 0 | | |
| Acar-UA*144 | 1 | | |
| Acar-UA*188 | 0 | | |
| Acar-UA*243 | 0 | | |
| Acar-UA*304 | 0 | | |

| **Acar-HPLT*76** | | Prop. unresolved alleles | 0.166667 |
|---|---|---|---|
| Allele | Unresolved | | |
| Acar-UA*4 | 0 | | |
| Acar-UA*9 | 1 | | |
| Acar-UA*30 | 0 | | |
| Acar-UA*55 | 0 | | |
| Acar-UA*144 | 0 | | |
| Acar-UA*242 | 0 | | |

| **Acar-HPLT*77** | | Prop. unresolved alleles | 0.444444 |
|---|---|---|---|
| Allele | Unresolved | | |
| Acar-UA*4 | 1 | | |
| Acar-UA*9 | 1 | | |
| Acar-UA*48 | 0 | | |
| Acar-UA*55 | 1 | | |
| Acar-UA*122 | 0 | | |
| Acar-UA*144 | 1 | | |
| Acar-UA*162 | 0 | | |
| Acar-UA*174 | 0 | | |
| Acar-UA*277 | 0 | | |

| **Acar-HPLT*78** | | Prop. unresolved alleles | 0 |
|---|---|---|---|
| Allele | Unresolved | | |
| Acar-UA*12 | 0 | | |
| Acar-UA*50 | 0 | | |
| Acar-UA*55 | 0 | | |
| Acar-UA*181 | 0 | | |
| Acar-UA*208 | 0 | | |
| Acar-UA*248 | 0 | | |
| Acar-UA*373 | 0 | | |

| **Acar-HPLT*79** | | Prop. unresolved alleles | 0.4 |
|---|---|---|---|
| Allele | Unresolved | | |
| Acar-UA*4 | 1 | | |
| Acar-UA*9 | 1 | | |
| Acar-UA*55 | 1 | | |
| Acar-UA*71 | 0 | | |
| Acar-UA*137 | 0 | | |
| Acar-UA*144 | 1 | | |
| Acar-UA*250 | 0 | | |
| Acar-UA*279 | 0 | | |
| Acar-UA*322 | 0 | | |
| Acar-UA*328 | 0 | | |

| **Acar-HPLT*80** | | Prop. unresolved alleles | 0.230769 |
|---|---|---|---|
| Allele | Unresolved | | |
| Acar-UA*4 | 1 | | |
| Acar-UA*9 | 1 | | |
| Acar-UA*51 | 0 | | |
| Acar-UA*52 | 0 | | |
| Acar-UA*55 | 1 | | |
| Acar-UA*107 | 0 | | |
| Acar-UA*110 | 0 | | |
| Acar-UA*143 | 0 | | |
| Acar-UA*144 | 0 | | |
| Acar-UA*255 | 0 | | |
| Acar-UA*304 | 0 | | |
| Acar-UA*368 | 0 | | |
| Acar-UA*392 | 0 | | |

| **Acar-HPLT*81** | | Prop. unresolved alleles | 0.333333 |
|---|---|---|---|
| Allele | Unresolved | | |
| Acar-UA*4 | 1 | | |
| Acar-UA*9 | 1 | | |
| Acar-UA*11 | 0 | | |
| Acar-UA*47 | 0 | | |
| Acar-UA*55 | 1 | | |
| Acar-UA*128 | 0 | | |
| Acar-UA*144 | 0 | | |
| Acar-UA*150 | 0 | | |
| Acar-UA*332 | 0 | | |

| **Acar-HPLT*82** | | Prop. unresolved alleles | 0.75 |
|---|---|---|---|
| Allele | Unresolved | | |
| Acar-UA*4 | 1 | | |
| Acar-UA*9 | 1 | | |
| Acar-UA*55 | 1 | | |
| Acar-UA*254 | 0 | | |

| **Acar-HPLT*83** | | Prop. unresolved alleles | 0.333333 |
|---|---|---|---|
| Allele | Unresolved | | |
| Acar-UA*4 | 0 | | |
| Acar-UA*9 | 1 | | |
| Acaru-UA*23 | 0 | | |
| Acar-UA*55 | 1 | | |
| Acar-UA*285 | 0 | | |
| Acar-UA*342 | 0 | | |

| **Acar-HPLT*84** | | Prop. unresolved alleles | 0.095238 |
|---|---|---|---|
| Allele | Unresolved | | |
| Acar-UA*4 | 0 | | |
| Acar-UA*9 | 1 | | |
| Acar-UA*46 | 0 | | |
| Acar-UA*48 | 0 | | |
| Acar-UA*49 | 0 | | |
| Acar-UA*53 | 0 | | |
| Acar-UA*55 | 1 | | |
| Acar-UA*77 | 0 | | |
| Acar-UA*94 | 0 | | |
| Acar-UA*103 | 0 | | |
| Acar-UA*107 | 0 | | |
| Acar-UA*143 | 0 | | |
| Acar-UA*150 | 0 | | |
| Acar-UA*154 | 0 | | |
| Acar-UA*255 | 0 | | |
| Acar-UA*273 | 0 | | |
| Acar-UA*307 | 0 | | |
| Acar-UA*332 | 0 | | |
| Acar-UA*349 | 0 | | |
| Acar-UA*356 | 0 | | |
| Acar-UA*417 | 0 | | |

| **Acar-HPLT*85** | | Prop. unresolved alleles | 0.5 |
|---|---|---|---|
| Allele | Unresolved | | |
| Acar-UA*4 | 1 | | |
| Acar-UA*9 | 1 | | |
| Acar-UA*12 | 0 | | |
| Acar-UA*55 | 1 | | |
| Acar-UA*128 | 0 | | |
| Acar-UA*144 | 1 | | |
| Acar-UA*200 | 0 | | |
| Acar-UA*328 | 0 | | |

| **Acar-HPLT*86** | | Prop. unresolved alleles | 0.222222 |
|---|---|---|---|
| Allele | Unresolved | | |
| Acar-UA*9 | 1 | | |
| Acar-UA*33 | 0 | | |
| Acar-UA*55 | 1 | | |
| Acar-UA*94 | 0 | | |
| Acar-UA*144 | 0 | | |
| Acar-UA*151 | 0 | | |
| Acar-UA*341 | 0 | | |
| Acar-UA*364 | 0 | | |
| Acar-UA*404 | 0 | | |

| **Acar-HPLT*87** | | Prop. unresolved alleles | 0.333333 |
|---|---|---|---|
| Allele | Unresolved | | |
| Acar-UA*4 | 0 | | |
| Acar-UA*9 | 1 | | |
| Acaru-UA*23 | 0 | | |
| Acar-UA*55 | 1 | | |
| Acar-UA*144 | 0 | | |
| Acar-UA*285 | 0 | | |

| **Acar-HPLT*88** | | Prop. unresolved alleles | 0.25 |
|---|---|---|---|
| Allele | Unresolved | | |
| Acar-UA*4 | 1 | | |
| Acar-UA*9 | 1 | | |
| Acar-UA*12 | 0 | | |
| Acar-UA*48 | 0 | | |
| Acar-UA*55 | 1 | | |
| Acar-UA*96 | 0 | | |
| Acar-UA*145 | 0 | | |
| Acar-UA*173 | 0 | | |
| Acar-UA*214 | 0 | | |
| Acar-UA*279 | 0 | | |
| Acar-UA*304 | 0 | | |
| Acar-UA*330 | 0 | | |

| **Acar-HPLT*89** | | Prop. unresolved alleles | 0.384615 |
|---|---|---|---|
| Allele | Unresolved | | |
| Acar-UA*4 | 1 | | |
| Acar-UA*9 | 1 | | |
| Acar-UA*31 | 0 | | |
| Acar-UA*55 | 1 | | |
| Acar-UA*91 | 0 | | |
| Acar-UA*96 | 0 | | |
| Acar-UA*103 | 0 | | |
| Acar-UA*122 | 1 | | |
| Acar-UA*144 | 1 | | |
| Acar-UA*150 | 0 | | |
| Acar-UA*183 | 0 | | |
| Acar-UA*196 | 0 | | |
| Acar-UA*204 | 0 | | |

| **Acar-HPLT*90** | | Prop. unresolved alleles | 0.272727 |
|---|---|---|---|
| Allele | Unresolved | | |
| Acar-UA*4 | 0 | | |
| Acar-UA*8 | 0 | | |
| Acar-UA*9 | 1 | | |
| Acar-UA*12 | 1 | | |
| Acar-UA*33 | 0 | | |
| Acar-UA*42 | 0 | | |
| Acar-UA*55 | 1 | | |
| Acar-UA*144 | 0 | | |
| Acar-UA*156 | 0 | | |
| Acar-UA*163 | 0 | | |
| Acar-UA*248 | 0 | | |

| **Acar-HPLT*91** | | Prop. unresolved alleles | 0.363636 |
|---|---|---|---|
| Allele | Unresolved | | |
| Acar-UA*4 | 1 | | |
| Acar-UA*9 | 1 | | |
| Acar-UA*48 | 1 | | |
| Acar-UA*50 | 0 | | |
| Acar-UA*55 | 1 | | |
| Acar-UA*86 | 0 | | |
| Acar-UA*91 | 0 | | |
| Acar-UA*144 | 0 | | |
| Acar-UA*150 | 0 | | |
| Acar-UA*332 | 0 | | |
| Acar-UA*415 | 0 | | |

| **Acar-HPLT*92** | | Prop. unresolved alleles | 0.230769 |
|---|---|---|---|
| Allele | Unresolved | | |
| Acar-UA*9 | 1 | | |
| Acar-UA*55 | 1 | | |
| Acar-UA*126 | 0 | | |
| Acar-UA*144 | 1 | | |
| Acar-UA*150 | 0 | | |
| Acar-UA*208 | 0 | | |
| Acar-UA*229 | 0 | | |
| Acar-UA*328 | 0 | | |
| Acar-UA*349 | 0 | | |
| Acar-UA*380 | 0 | | |
| Acar-UA*407 | 0 | | |
| Acar-UA*412 | 0 | | |
| Acar-UA*414 | 0 | | |

| **Acar-HPLT*93** | | Prop. unresolved alleles | 0.555556 |
|---|---|---|---|
| Allele | Unresolved | | |
| Acar-UA*4 | 1 | | |
| Acar-UA*12 | 1 | | |
| Acar-UA*31 | 0 | | |
| Acar-UA*48 | 0 | | |
| Acar-UA*50 | 0 | | |
| Acar-UA*55 | 1 | | |
| Acar-UA*82 | 0 | | |
| Acar-UA*144 | 1 | | |
| Acar-UA*304 | 1 | | |

| **Acar-HPLT*94** | | Prop. unresolved alleles | 0.222222 |
|---|---|---|---|
| Allele | Unresolved | | |
| Acar-UA*9 | 1 | | |
| Acar-UA*11 | 0 | | |
| Acar-UA*47 | 0 | | |
| Acar-UA*55 | 1 | | |
| Acar-UA*128 | 0 | | |
| Acar-UA*141 | 0 | | |
| Acar-UA*144 | 0 | | |
| Acar-UA*150 | 0 | | |
| Acar-UA*332 | 0 | | |

| **Acar-HPLT*95** | | Prop. unresolved alleles | 0.166667 |
|---|---|---|---|
| Allele | Unresolved | | |
| Acar-UA*4 | 0 | | |
| Acar-UA*9 | 1 | | |
| Acar-UA*40 | 0 | | |
| Acar-UA*55 | 1 | | |
| Acar-UA*91 | 0 | | |
| Acar-UA*191 | 0 | | |
| Acar-UA*230 | 0 | | |
| Acar-UA*281 | 0 | | |
| Acar-UA*313 | 0 | | |
| Acar-UA*360 | 0 | | |
| Acar-UA*380 | 0 | | |
| Acar-UA*384 | 0 | | |

| **Acar-HPLT*96** | | Prop. unresolved alleles | 0.555556 |
|---|---|---|---|
| Allele | Unresolved | | |
| Acar-UA*4 | 1 | | |
| Acar-UA*9 | 1 | | |
| Acar-UA*11 | 0 | | |
| Acar-UA*12 | 1 | | |
| Acar-UA*31 | 0 | | |
| Acar-UA*55 | 1 | | |
| Acar-UA*144 | 1 | | |
| Acar-UA*183 | 0 | | |
| Acar-UA*387 | 0 | | |

| **Acar-HPLT*97** | | Prop. unresolved alleles | 0.25 |
|---|---|---|---|
| Allele | Unresolved | | |
| Acar-UA*9 | 1 | | |
| Acar-UA*55 | 1 | | |
| Acar-UA*128 | 0 | | |
| Acar-UA*144 | 0 | | |
| Acar-UA*191 | 0 | | |
| Acar-UA*337 | 0 | | |
| Acar-UA*371 | 0 | | |
| Acar-UA*410 | 0 | | |

| **Acar-HPLT*98** | | Prop. unresolved alleles | 0.5 |
|---|---|---|---|
| Allele | Unresolved | | |
| Acar-UA*9 | 1 | | |
| Acar-UA*55 | 1 | | |
| Acar-UA*144 | 1 | | |
| Acar-UA*210 | 0 | | |
| Acar-UA*229 | 0 | | |
| Acar-UA*375 | 0 | | |

| **Acar-HPLT*99** | | Prop. unresolved alleles | 0.333333 |
|---|---|---|---|
| Allele | Unresolved | | |
| Acar-UA*4 | 1 | | |
| Acar-UA*9 | 1 | | |
| Acar-UA*48 | 0 | | |
| Acar-UA*50 | 0 | | |
| Acar-UA*55 | 1 | | |
| Acar-UA*90 | 0 | | |
| Acar-UA*144 | 0 | | |
| Acar-UA*146 | 1 | | |
| Acar-UA*216 | 0 | | |
| Acar-UA*265 | 0 | | |
| Acar-UA*266 | 0 | | |
| Acar-UA*378 | 0 | | |

| **Acar-HPLT*100** | | Prop. unresolved alleles | 0.3 |
|---|---|---|---|
| Allele | Unresolved | | |
| Acar-UA*9 | 1 | | |
| Acaru-UA*23 | 0 | | |
| Acar-UA*41 | 0 | | |
| Acar-UA*55 | 1 | | |
| Acar-UA*94 | 1 | | |
| Acar-UA*103 | 0 | | |
| Acar-UA*144 | 0 | | |
| Acar-UA*192 | 0 | | |
| Acar-UA*200 | 0 | | |
| Acar-UA*342 | 0 | | |

| **Acar-HPLT*101** | | Prop. unresolved alleles | 0.444444 |
|---|---|---|---|
| Allele | Unresolved | | |
| Acar-UA*4 | 1 | | |
| Acar-UA*9 | 1 | | |
| Acar-UA*55 | 1 | | |
| Acar-UA*66 | 0 | | |
| Acar-UA*111 | 0 | | |
| Acar-UA*144 | 1 | | |
| Acar-UA*181 | 0 | | |
| Acar-UA*406 | 0 | | |
| Acar-UA*411 | 0 | | |

| **Acar-HPLT*102** | | Prop. unresolved alleles | 0.363636 |
|---|---|---|---|
| Allele | Unresolved | | |
| Acar-UA*4 | 1 | | |
| Acar-UA*9 | 1 | | |
| Acar-UA*55 | 1 | | |
| Acar-UA*94 | 0 | | |
| Acar-UA*122 | 0 | | |
| Acar-UA*125 | 0 | | |
| Acar-UA*144 | 1 | | |
| Acar-UA*304 | 0 | | |
| Acar-UA*405 | 0 | | |
| Acar-UA*409 | 0 | | |
| Acar-UA*413 | 0 | | |

| **Acar-HPLT*103** | | Prop. unresolved alleles | 0.333333 |
|---|---|---|---|
| Allele | Unresolved | | |
| Acar-UA*4 | 1 | | |
| Acar-UA*6 | 0 | | |
| Acar-UA*8 | 0 | | |
| Acar-UA*9 | 1 | | |
| Acar-UA*11 | 0 | | |
| Acar-UA*55 | 0 | | |
| Acar-UA*73 | 0 | | |
| Acar-UA*144 | 1 | | |
| Acar-UA*156 | 0 | | |

| **Acar-HPLT*104** | | Prop. unresolved alleles | 0.222222 |
|---|---|---|---|
| Allele | Unresolved | | |
| Acar-UA*4 | 1 | | |
| Acar-UA*9 | 0 | | |
| Acar-UA*31 | 0 | | |
| Acar-UA*48 | 0 | | |
| Acar-UA*109 | 0 | | |
| Acar-UA*144 | 1 | | |
| Acar-UA*241 | 0 | | |
| Acar-UA*330 | 0 | | |
| Acar-UA*400 | 0 | | |

| **Acar-HPLT*105** | | Prop. unresolved alleles | 0.5 |
|---|---|---|---|
| Allele | Unresolved | | |
| Acar-UA*4 | 1 | | |
| Acar-UA*9 | 1 | | |
| Acar-UA*55 | 1 | | |
| Acar-UA*144 | 1 | | |
| Acar-UA*191 | 0 | | |
| Acar-UA*337 | 0 | | |
| Acar-UA*371 | 0 | | |
| Acar-UA*410 | 0 | | |

| **Acar-HPLT*106** | | Prop. unresolved alleles | 0.25 |
|---|---|---|---|
| Allele | Unresolved | | |
| Acar-UA*4 | 1 | | |
| Acar-UA*9 | 1 | | |
| Acar-UA*40 | 0 | | |
| Acar-UA*55 | 1 | | |
| Acar-UA*91 | 0 | | |
| Acar-UA*144 | 0 | | |
| Acar-UA*191 | 0 | | |
| Acar-UA*281 | 0 | | |
| Acar-UA*290 | 0 | | |
| Acar-UA*310 | 0 | | |
| Acar-UA*380 | 0 | | |
| Acar-UA*384 | 0 | | |

| **Acar-HPLT*107** | | Prop. unresolved alleles | 0.4 |
|---|---|---|---|
| Allele | Unresolved | | |
| Acar-UA*4 | 1 | | |
| Acar-UA*9 | 1 | | |
| Acar-UA*33 | 0 | | |
| Acar-UA*48 | 1 | | |
| Acar-UA*55 | 1 | | |
| Acar-UA*122 | 0 | | |
| Acar-UA*144 | 0 | | |
| Acar-UA*163 | 0 | | |
| Acar-UA*277 | 0 | | |
| Acar-UA*384 | 0 | | |

### Pedigree table

| Breeding individual | Breeding year 1991 | 1996 | 1998 | 1999 | Ancestors Mother | Father | M GM | M GF | P GM | P GF | MM GGM | MM GGF | MP GGM | MP GGF | PM GGM | PM GGF | PP GGM | PP GGF | MMM GGGM | MMM GGGF | MMP GGGM | MMP GGGF | MPP GGGM | MPP GGGF | PMM GGGM | PMM GGGF | PMP GGGM | PMP GGGF | PPM GGGM | PPM GGGF | PPP GGGM | PPP GGGF | PPPP GGGGM | PPPP GGGGF |
|---|---|---|---|---|---|---|---|---|---|---|---|---|---|---|---|---|---|---|---|---|---|---|---|---|---|---|---|---|---|---|---|---|---|---|
| 289_ad | | | | x | | | | | | | | | | | | | | | | | | | | | | | | | | | | | | |
| 295_ad | | | | x | | | | | | | | | | | | | | | | | | | | | | | | | | | | | | |
| 296_ad | | | x | x | | | | | | | | | | | | | | | | | | | | | | | | | | | | | | |
| 297_ad | | | x | x | | | | | | | | | | | | | | | | | | | | | | | | | | | | | | |
| 304_ad | | | x | | | | | | | | | | | | | | | | | | | | | | | | | | | | | | | |
| 305_ad | | | | x | | | | | | | | | | | | | | | | | | | | | | | | | | | | | | |
| 307_ad | | | | x | | | | | | | | | | | | | | | | | | | | | | | | | | | | | | |
| 308_ad | | | | x | | | | | | | | | | | | | | | | | | | | | | | | | | | | | | |
| 309_ad | | | | x | | | | | | | | | | | | | | | | | | | | | | | | | | | | | | |
| 310_ad | | | | x | | | | | | | | | | | | | | | | | | | | | | | | | | | | | | |
| 311_ad | | | | x | | | | | | | | | | | | | | | | | | | | | | | | | | | | | | |
| 312_ad | | | | x | | | | | | | | | | | | | | | | | | | | | | | | | | | | | | |
| 319_ad | x | | | | | | | | | | | | | | | | | | | | | | | | | | | | | | | | | |
| 332_ad | | x | | | 319_ad | 466_ad | | | 560_ad | 561_ad | | | | | | | | | | | | | | | | | | | | | | | | |
| 341_ad | | | x | x | 391_ad | 412_ad | 430_ad | 513_ad | | | 315_ad | 323_ad | | | | | | | | | | | | | | | | | | | | | | |
| 361_ad | | | x | | 338_ad | 422_ad | 502_ad | 438_ad | | | | | | | | | | | | | | | | | | | | | | | | | | |
| 362_ad | | | | x | 391_ad | 422_ad | 430_ad | 513_ad | | | 315_ad | 323_ad | | | | | | | | | | | | | | | | | | | | | | |
| 363_ad | | | | x | 334_ad | 438_ad | 313_ad | 360_ad | | | | | 411_ad | 428_ad | | | | | | | | | | 326_ad | | | | | | | | | | |
| 366_ad | | | x | x | 424_ad | 465_ad | | | 488_ad | 432_ad | | | | | | | 319_ad | 562_ad | | | | | | | | | | | | | | | | |
| 370_ad | | | x | | 532_ad | 365_ad | | | 334_ad | 438_ad | | | | | 313_ad | 360_ad | | | | | | | | | | | 411_ad | 428_ad | | | | | | |
| 371_ad | | | x | x | 351_ad | 425_ad | 339_ad | 442_ad | | | 433_ad | 384_ad | | | | | | | | | 315_ad | 326_ad | | | | | | | | | | | | |
| 373_ad | | | x | | 294_ad | 436_ad | | | | | | | | | | | | | | | | | | | | | | | | | | | | |
| 375_ad | | | x | | 558_ad | 521_ad | | | | | | | | | | | | | | | | | | | | | | | | | | | | |
| 376_ad | | | x | x | 294_ad | 545_ad | | | | | | | | | | | | | | | | | | | | | | | | | | | | |
| 377_ad | | | x | | 538_ad | 542_ad | | | | | | | | | | | | | | | | | | | | | | | | | | | | |
| 378_ad | | | x | | 342_ad | 357_ad | 329_ad | 418_ad | 441_ad | 398_ad | 317_ad | 438_ad | 315_ad | 323_ad | | | 313_ad | 561_ad | | | | | | | | | | | | | | | | |
| 380_ad | | | x | | 424_ad | 546_ad | | | | | | | | | | | | | | | | | | | | | | | | | | | | |
| 385_ad | | | x | x | 403_ad | 546_ad | | | | | | | | | | | | | | | | | | | | | | | | | | | | |
| 386_ad | | | x | | 407_ad | 425_ad | 476_ad | 332_ad | | | 313_ad | 428_ad | 319_ad | 466_ad | | | | | | | | 326_ad | 560_ad | 561_ad | | | | | | | | | | |
| 388_ad | | | x | x | 522_ad | 357_ad | | | 441_ad | 398_ad | | | | | | | 313_ad | 561_ad | | | | | | | | | | | | | | | | |
| 390_ad | | | x | x | 346_ad | 355_ad | 338_ad | 454_ad | 478_ad | 347_ad | 502_ad | 438_ad | 313_ad | 561_ad | 379_ad | 396_ad | 435_ad | 418_ad | | | | | | | 433_ad | | | 514_ad | | | 315_ad | 323_ad | | |
| 394_ad | | | | x | 391_ad | 340_ad | 430_ad | 513_ad | 391_ad | 412_ad | 315_ad | 323_ad | | | 430_ad | 513_ad | | | | | | | | | 315_ad | 323_ad | | | | | | | | |
| 395_ad | | | x | x | 403_ad | 546_ad | | | | | | | | | | | | | | | | | | | | | | | | | | | | |
| 401_ad | | | x | x | 533_ad | 546_ad | | | | | | | | | | | | | | | | | | | | | | | | | | | | |
| 403_ad | | | x | | | | | | | | | | | | | | | | | | | | | | | | | | | | | | | |
| 414_ad | | | x | | | | | | | | | | | | | | | | | | | | | | | | | | | | | | | |
| 419_ad | | | | x | | | | | | | | | | | | | | | | | | | | | | | | | | | | | | |
| 424_ad | | | x | | | | | | | | | | | | | | | | | | | | | | | | | | | | | | | |
| 426_ad | | | | x | | | | | | | | | | | | | | | | | | | | | | | | | | | | | | |
| 448_ad | | | | x | 424_ad | 361_ad | | | 338_ad | 422_ad | | | | | 502_ad | 438_ad | | | | | | | | | | | | | | | | | | |
| 449_ad | | | | x | 539_ad | 536_ad | | | | | | | | | | | | | | | | | | | | | | | | | | | | |
| 450_ad | | | x | x | 290_ad | 296_ad | | | | | | | | | | | | | | | | | | | | | | | | | | | | |
| 451_ad | | | | x | 403_ad | 355_ad | | | 478_ad | 347_ad | | | | | 379_ad | 396_ad | 435_ad | 418_ad | | | | | | | 433_ad | | | 514_ad | | | 315_ad | 323_ad | | |
| 453_ad | | | x | x | 539_ad | 536_ad | | | | | | | | | | | | | | | | | | | | | | | | | | | | |
| 456_ad | | | | x | 426_ad | 340_ad | | | 391_ad | 412_ad | | | | | 430_ad | 513_ad | | | | | | | | | 315_ad | 323_ad | | | | | | | | |
| 457_ad | | | x | | 288_ad | 516_ad | | | | | | | | | | | | | | | | | | | | | | | | | | | | |
| 458_ad | | | | x | 539_ad | 536_ad | | | | | | | | | | | | | | | | | | | | | | | | | | | | |
| 460_ad | | | x | x | 362_ad | 516_ad | 391_ad | 422_ad | | | 430_ad | 513_ad | | | | | | | 315_ad | 323_ad | | | | | | | | | | | | | | |
| 462_ad | | | | x | 527_ad | 355_ad | | | 478_ad | 347_ad | | | | | 379_ad | 396_ad | 435_ad | 418_ad | | | | | | | | | | 514_ad | | | 315_ad | 323_ad | | |
| 464_ad | | | x | x | 341_ad | 340_ad | 391_ad | 412_ad | 391_ad | 412_ad | 430_ad | 513_ad | | | 430_ad | 513_ad | | | 315_ad | 323_ad | | | | | 315_ad | 323_ad | | | | | | | | |
| 466_ad | x | | | | 560_ad | 561_ad | | | | | | | | | | | | | | | | | | | | | | | | | | | | |
| 467_ad | | | x | x | 541_ad | 525_ad | | | | | | | | | | | | | | | | | | | | | | | | | | | | |
| 468_ad | | | | x | 366_ad | 390_ad | 424_ad | 465_ad | 346_ad | 355_ad | | | 488_ad | 432_ad | 338_ad | 454_ad | 478_ad | 347_ad | | | | | 319_ad | 562_ad | 502_ad | 438_ad | 313_ad | 561_ad | 379_ad | 396_ad | 435_ad | 418_ad | 315_ad | 323_ad |
| 471_ad | | | | x | 457_ad | 524_ad | 288_ad | 516_ad | | | | | | | | | | | | | | | | | | | | | | | | | | |
| 479_ad | | | | x | 373_ad | 385_ad | 294_ad | 436_ad | 403_ad | 546_ad | | | | | | | | | | | | | | | | | | | | | | | | |
| 481_ad | | | | x | | | | | | | | | | | | | | | | | | | | | | | | | | | | | | |
| 516_ad | | | x | x | | | | | | | | | | | | | | | | | | | | | | | | | | | | | | |
| 518_ad | | | x | x | | | | | | | | | | | | | | | | | | | | | | | | | | | | | | |
| 519_ad | | | x | x | | | | | | | | | | | | | | | | | | | | | | | | | | | | | | |
| 520_ad | | | x | x | | | | | | | | | | | | | | | | | | | | | | | | | | | | | | |
| 523_ad | | | x | | | | | | | | | | | | | | | | | | | | | | | | | | | | | | | |
| 524_ad | | | x | | | | | | | | | | | | | | | | | | | | | | | | | | | | | | | |
| 527_ad | | | x | | | | | | | | | | | | | | | | | | | | | | | | | | | | | | | |
| 529_ad | | | x | x | | | | | | | | | | | | | | | | | | | | | | | | | | | | | | |
| 530_ad | | | x | | | | | | | | | | | | | | | | | | | | | | | | | | | | | | | |
| 531_ad | | | x | x | | | | | | | | | | | | | | | | | | | | | | | | | | | | | | |
| 534_ad | | | | x | | | | | | | | | | | | | | | | | | | | | | | | | | | | | | |
| 535_ad | | | x | x | | | | | | | | | | | | | | | | | | | | | | | | | | | | | | |
| 537_ad | | | | x | | | | | | | | | | | | | | | | | | | | | | | | | | | | | | |
| 538_ad | | | x | | | | | | | | | | | | | | | | | | | | | | | | | | | | | | | |
| 539_ad | | | x | x | | | | | | | | | | | | | | | | | | | | | | | | | | | | | | |
| 541_ad | | | x | | | | | | | | | | | | | | | | | | | | | | | | | | | | | | | |
| 543_ad | | | x | | | | | | | | | | | | | | | | | | | | | | | | | | | | | | | |
| 544_ad | | | x | | | | | | | | | | | | | | | | | | | | | | | | | | | | | | | |
| 547_ad | | | | x | | | | | | | | | | | | | | | | | | | | | | | | | | | | | | |
| 549_ad | | x | | | | | | | | | | | | | | | | | | | | | | | | | | | | | | | | |
| 550_ad | | | x | x | | | | | | | | | | | | | | | | | | | | | | | | | | | | | | |
| 551_ad | | | x | | | | | | | | | | | | | | | | | | | | | | | | | | | | | | | |
| 552_ad | | | | x | | | | | | | | | | | | | | | | | | | | | | | | | | | | | | |
| 554_ad | | | x | | | | | | | | | | | | | | | | | | | | | | | | | | | | | | | |
| 555_ad | | | | x | | | | | | | | | | | | | | | | | | | | | | | | | | | | | | |
| 557_ad | | | | x | | | | | | | | | | | | | | | | | | | | | | | | | | | | | | |

Legend

| | |
|---|---|
| M GM | Maternal (mat) grandmother |
| M GF | Mat grandfather |
| P GM | Paternal (pat) grandmother |
| P GF | Pat grandfather |
| MM GGM | Mat mat great grandmother |
| MM GGF | Mat mat great grandfather |
| MP GGM | Mat pat great grandmother |
| MP GGF | Mat pat great grandfather |
| PM GGM | Pat mat great grandmother |
| PM GGF | Pat mat great grandfather |
| PP GGM | Pat pat great grandmother |
| PP GGF | Pat pat great grandfather |
| MMM GGGM | Mat mat mat great great grandmother |
| MMM GGGF | Mat mat mat great great grandfather |
| MMP GGGM | Mat mat pat great great grandmother |
| MMP GGGF | Mat mat pat great great grandfather |
| MPP GGGM | Mat pat pat great great grandmother |
| MPP GGGF | Mat pat pat great great grandfather |
| PMM GGGM | Pat mat mat great great grandmother |
| PMM GGGF | Pat mat mat great great grandfather |
| PMP GGGM | Pat mat pat great great grandmother |
| PMP GGGF | Pat mat pat great great grandfather |
| PPM GGGM | Pat pat mat great great grandmother |
| PPM GGGF | Pat pat mat great great grandfather |
| PPP GGGM | Pat pat pat great great grandmother |
| PPP GGGF | Pat pat pat great great grandfather |
| PPPP GGGGM | Pat pat pat pat great great great grandmother |
| PPPP GGGGF | Pat pat pat pat great great great grandfather |

The individual codes given in the table correspond to the unique color codes that the birds were given when ringed. Light and dark gray background shading indicate maternal and paternal ancestral lineages, respectively.

Three ancestral individuals (560_ad, 561_ad, and 562_ad) were not genotyped, and thus not included in the data set. These are marked with a streaked background pattern in the pedigree table.
